# Glioma Cells Secrete Collagen VI to Facilitate Invasion

**DOI:** 10.1101/2023.12.12.571198

**Authors:** Junghwa Cha, Erika A. Ding, Emily M. Carvalho, Annabelle Fowler, Manish K. Aghi, Sanjay Kumar

## Abstract

While glioblastoma (GBM) progression is associated with extensive extracellular matrix (ECM) secretion, the causal contributions of ECM secretion to invasion remain unclear. Here we investigate these contributions by combining engineered materials, proteomics, analysis of patient data, and a model of bevacizumab-resistant GBM. We find that GBM cells cultured in engineered 3D hyaluronic acid hydrogels secrete ECM prior to invasion, particularly in the absence of exogenous ECM ligands. Proteomic measurements reveal extensive secretion of collagen VI, and collagen VI-associated transcripts are correspondingly enriched in microvascular proliferation regions of human GBMs. We further show that bevacizumab-resistant GBM cells deposit more collagen VI than their responsive counterparts, which is associated with marked cell-ECM stiffening. COL6A3 deletion in GBM cells reduces invasion, β-catenin signaling, and expression of mesenchymal markers, and these effects are amplified in hypoxia. Our studies strongly implicate GBM cell-derived collagen VI in microenvironmental remodeling to facilitate invasion.

## Introduction

Glioblastoma (GBM) is a rapidly progressive primary brain tumor characterized by extensive infiltration of brain tissue by tumor cells (*1*). GBMs dramatically alter the normal brain extracellular matrix (ECM) to create a pro-invasive microenvironment (*2*), although the details of this remodeling process remain incompletely understood. Normal brain tissue is dominated by a complex mixture of high molecular weight hyaluronic acid (HA), proteoglycans, and glycoproteins (*3, 4*). Unlike connective tissues, fibrous ECM components are largely absent and traditionally viewed as restricted to perivascular spaces (*5*). Among solid tissues, brain tissue is also exceptionally soft, with elastic moduli typically measured in the <1 kPa range (*6–9*). During GBM progression, tumor and stromal cells increase deposition of many of these components, including collagens (*10*), fibronectin (*11*), and tenascin-C (*12*), and HA is both more abundant and shifted towards lower molecular weight distributions (*13*). Tumor angiogenesis also contributes to the upregulation of ECM components to support greater vascularization, and the deposition of ECMs around blood vessels can itself drive directional cell migration (*14*). Because of these ECM changes and the increased cellularity of the tumor, GBM tissue can become dramatically stiffer than normal brain (*15*), with a recent study reporting compressive moduli of 1-10 kPa in the tumor core and edge (*16, 17*).

Changes in ECM composition and mechanics functionally contribute to tumor invasion on several levels. First, the spatially heterogeneous composition and structure of tumor ECM contribute to the stereotypical invasion patterns of GBM (*18*), which preferentially follow anatomical “seams” such as white matter tracts, perivascular beds, and the corpus callosum (*7, 19, 20*). Many glycoproteins enriched in the GBM microenvironment, such as osteopontin and tenascin-C, directly stimulate tumor cell motility and invasion (*21, 22*), and the abundance of low molecular weight HA species can signal through cellular HA receptors to promote GBM stemness and chemoresistance (*13, 23*). Chemoresistance responses, most notably to the anti-angiogenic agent bevacizumab, reciprocally induce ECM remodeling (*24*) while driving the mesenchymal transition of GBM and speeding invasion along perivascular tracts (*25*), following routes along the basement lamina of capillaries containing integrin ligands laminin, fibronectin, and collagen type IV. This perivascular invasion gives tumor cells access to the perivascular space whose proximity to vessels provides the oxygen, glucose, and nutrient-rich microenvironment needed for the survival of tumor cells and enables spontaneous, VEGF-independent tumor vascularization not affected by bevacizumab by engulfing preexisting brain microvessels. Finally, changes in ECM mechanics are increasingly believed to contribute to tumor infiltration, with higher ECM stiffness driving migration and proliferation in culture (*26*) and a variety of preclinical models (*27*).

Despite the clear functional significance of ECM remodeling to GBM invasion, angiogenesis, and chemoresistance, the field lacks insight into the earliest stages of matrix remodeling, when an initially normal ECM is converted into a pro-invasive microenvironment. Gaining a clearer picture of these early matrix secretion events would add to our mechanistic understanding of GBM invasion while also potentially exposing new therapeutic targets. However, inferring early events from *in vivo* models and patient data has proven challenging, in part because the tissue is typically harvested and analyzed only after the invasion is well underway. We and others have applied engineered 3D HA ECMs for *in vitro* modeling of GBM initiation and invasion (*19, 28–31*), as these materials recapitulate defining features of GBM invasion while permitting high-resolution imaging and analysis throughout the process.

In this study, we applied an engineered 3D HA ECM platform to investigate early ECM protein deposition events associated with invasion. Through a combination of metabolic labeling of secreted proteins, unbiased matrisomal analysis, and focused candidate-based studies, we identify collagen VI as a tumor-secreted driver of invasion. We further show that collagen VI stiffens the initially soft HA matrix and triggers mechanotransductive signaling through specific integrin subtypes to stimulate β-catenin signaling and expression of the mesenchymal protein ZEB1, the collective effect of which is to speed invasion. Collagen VI secretion is also greatly enhanced in a culture model of bevacizumab resistance, with suppression of collagen VI greatly attenuating hypoxia-driven invasion and expression of mesenchymal markers.

## Results

### GBM cells secrete protein and stiffen their surroundings following encapsulation in soft 3D HA hydrogels

Our first goal was to investigate how GBM cells remodel the surrounding ECM to facilitate invasion. As a model ECM platform, we used 3D engineered hyaluronic acid (HA) hydrogels, which we have used in the past due to their biochemical and biophysical resemblance to brain ECM (*28, 30, 32*). To provide a non-integrin adhesive background, we omitted RGD-based peptide ligands that are often used in these and other biomaterials to support integrin-mediated cell adhesion (hereafter referred to as HA/RGD-). We chose hydrogels with an elasticity of ∼250 Pa to approximate the stiffness of normal brain tissue (fig. S1A). Shortly after hydrogel encapsulation as single cells, U87 human glioma cells extended micro-protrusions, reflecting active engagement with the surrounding matrix (**Fig. 1A**). Over the course of 7 days, cells grew into multicellular aggregates with increasingly irregular borders driven by micro-protrusions, as evidenced by reduced shape factor by 7 days post-implantation (**Fig. 1B, C**). This contrasts with cells embedded within integrin-ligating matrices such as collagen I and Matrigel, where cells elongated within 3 days and largely remained identifiable as single cells (fig. S2).

**Figure 1.**
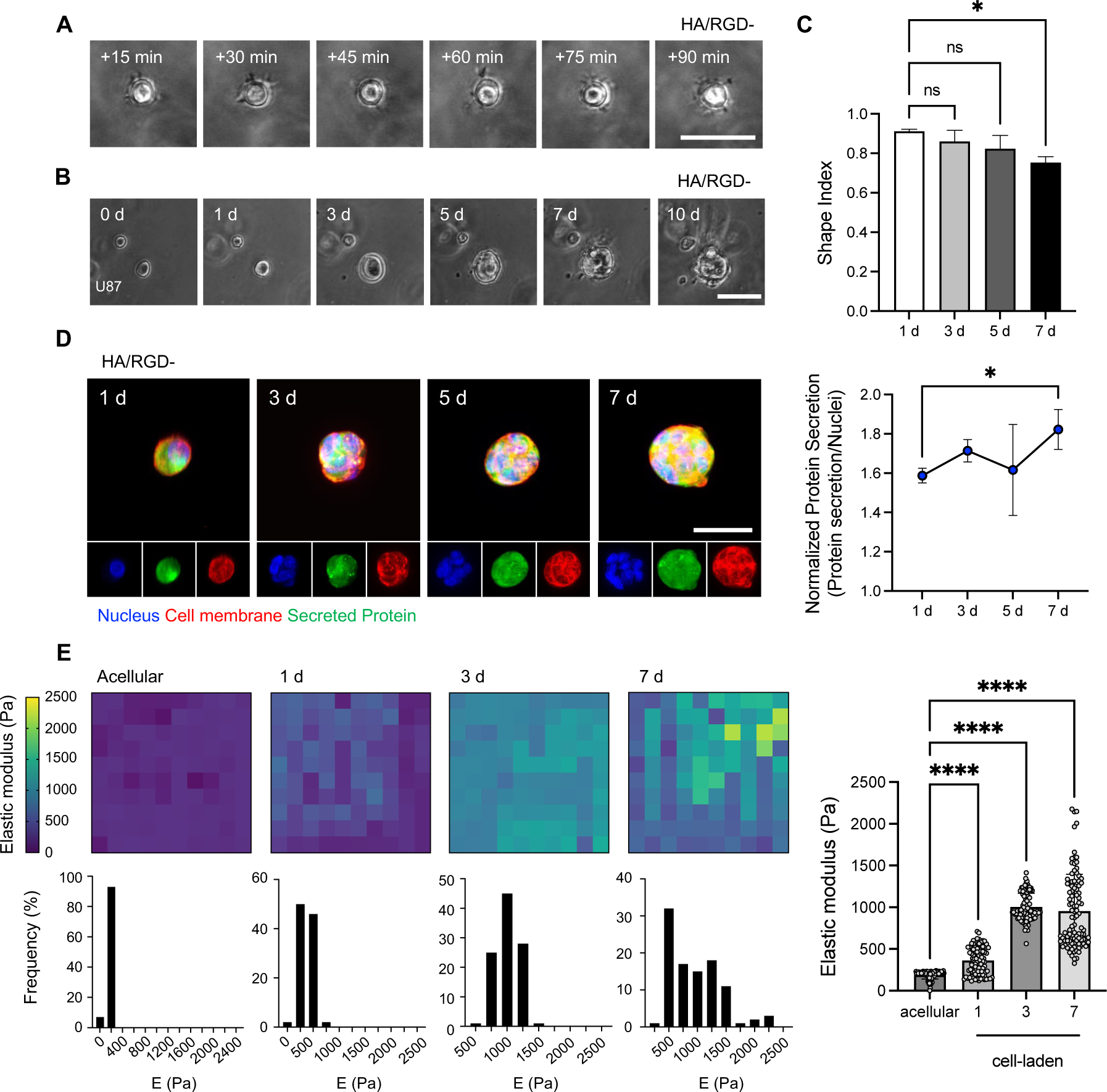
GBM cells remodel 3D HA hydrogels by secreting proteins. **(A)** Representative short­term time-lapse images of single U87 GBM cells interacting with HA/RGD-gels. Scale bar: 50 pm. (**B**) Representative time-lapse images of GBM cells encapsulated in HA/RGD-gels for 10 days. Scale bar: 50 pm. (**C**) Evolution of GBM cell shape index after HA/RGD-encapsulation. (n = 4 to 6) (**D**) Representative images of proteins secreted by GBM cells over time via metabolic labeling (left). Scale bar: 50 pm. Quantification of fluorescence signal from secreted proteins, normalized by the intensities of nuclei (right). (n = 4 to 6) (**E**) Representative force maps (top row) and histograms (bottom row) of the stiffness of acellular and cell-laden HA/RGD-gels over time (left). Comparison of matrix stiffness for acellular and cell-laden HA/RGD-gels over time (right). Statistical significance was analyzed using one-way ANOVA followed by Tukey’s multiple comparisons test, *p < 0.05, ****p < 0.001, ns: no significance.

Because HA/RGD-matrices lack endogenous integrin-ligating capabilities, we reasoned that cells might be interacting with these matrices through the secretion of ECM components. To test this hypothesis, we utilized a metabolic labeling technique to visualize cell-secreted proteins, including ECM proteins. This approach is based on supplementation of the culture medium with an azide-functionalized methionine analogue (azidohomoalanine, AHA), which is taken up and incorporated into newly synthesized proteins, enabling subsequent detection via bio-orthogonal click chemistry with a dibenzocyclooctyne (DBCO) tagged with a fluorophore (*33*). This metabolic labeling technique revealed extensive protein secretion starting as early as 1 day post-encapsulation and increasing over 7 days (**Fig. 1D**). Interestingly, the effect strongly depended on matrix stiffness, because when we increased the matrix stiffness of HA/RGD-hydrogels to 2.5 kPa (fig. S1B), approximating the stiffness of GBM tissue, cells formed spherical aggregates with well-defined boundaries and rarely developed micro-protrusions. Cells in stiff matrices also showed reduced overall protein secretion with little variation over 7 days (fig. S3). Intriguingly, protein secretion in soft HA/RGD-hydrogels increased microenvironmental stiffness by 5-fold over 7 days, as evidenced by atomic force microscopy (AFM) mapping of cell-laden hydrogels (**Fig. 1E**). Elasticity histograms additionally revealed greater mechanical heterogeneity with longer culture times.

### Protein deposition depends on matrix stiffness, integrin ligation, and protease degradability

Having established that GBM cells can engage 3D RGD-free ECMs of brain-like stiffness in a manner that is accompanied by protein secretion, we next asked whether these behaviors might differ in HA matrices that include pre-existing integrin ligands. We therefore repeated our experiments in soft (∼250 Pa) methacrylated HA hydrogels conjugated with RGD-containing peptides (Ac-GCGYGRGDSPG-NH2). In strong contrast to HA/RGD-matrices, protein deposition in HA/RGD+ matrices over 7 days was limited despite being accompanied by extended matrix protrusions (**Fig. 2A**, fig. S4A). However, protein deposition remained strongly stiffness-dependent, with stiffer (2.5 kPa) HA matrices supporting less ECM deposition than corresponding soft, RGD+ conjugated matrices.

**Figure 2.**
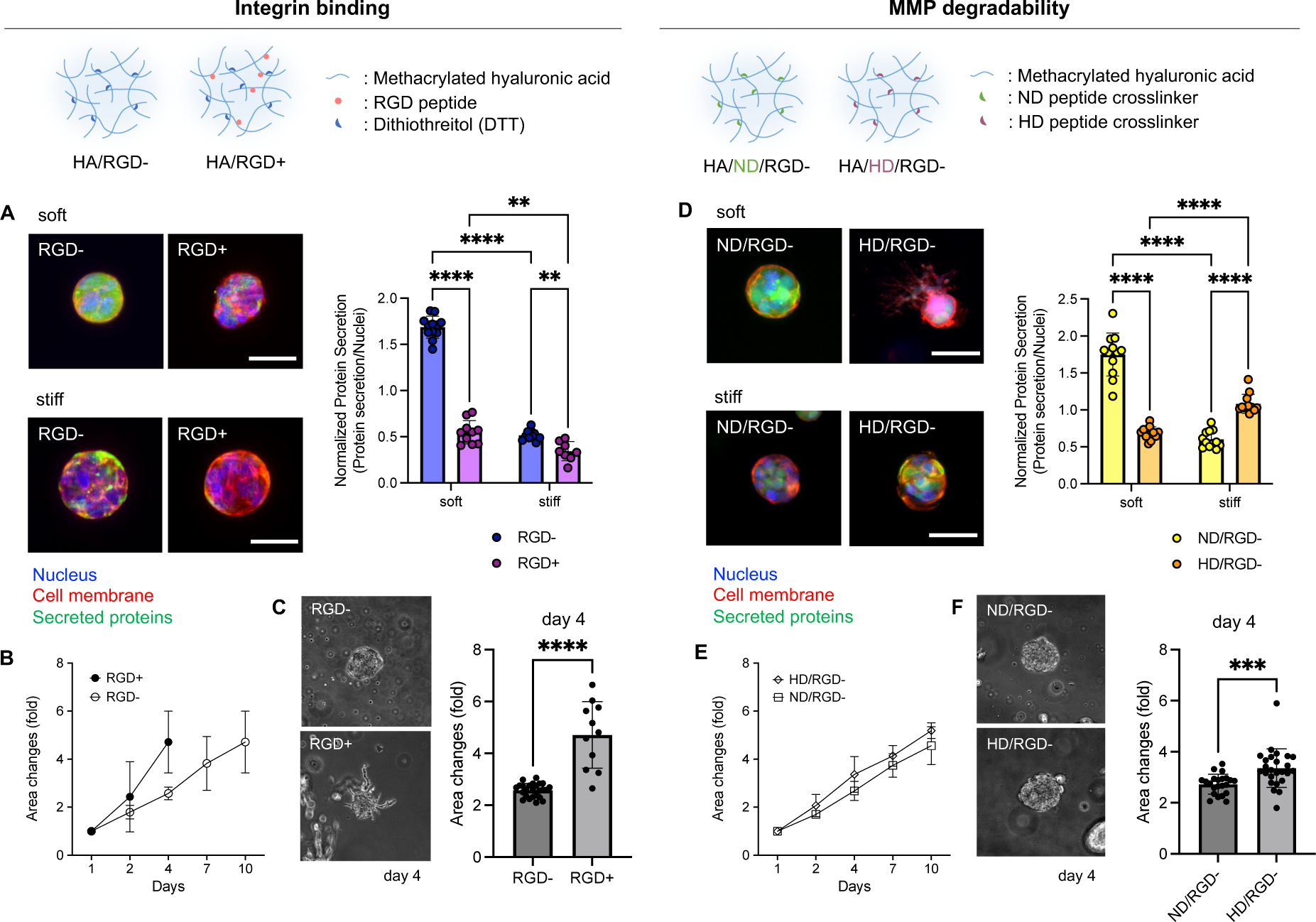
ECM remodeling by GBM depends on HA integrin ligation status (A-C) and protease degradability (D-F) (**A**) Representative images of fluorescently labeled secreted proteins by GBM cells depending on integrin binding (RGD-/+) (left). Scale bar: 50 μm. Quantification of fluorescence signals from secreted proteins by GBM cells, normalized by the intensities of nuclei. (right) (n = 10 to 12) (**B**) Quantification of 3D GBM tumorsphere invasion assay depending on integrin binding in the course of time up to 10 days. (n = 11 to 25) (**C**) Representative images of GBM tumorsphere invasion at day 4 within HA depending on integrin binding (RGD-/+) (left). Quantification of area changes at day 4 compared to day 0 (right). (**D**) Representative images of fluorescently labeled secreted proteins by GBM cells as a function of degradability (ND/RGD- and HD/RGD-) (left). ND: non-degradable crosslinker, HD: highly degradable crosslinker. Scale bar: 50 μm. Quantification of fluorescent signals from secreted proteins by GBM cells, normalized by the intensities of nuclei. (right) (n = 12) (**E**) Quantification of 3D GBM tumorsphere invasion assay depending on gel degradability in the course of time up to 10 days. (n = 22 to 25) (**F**) Representative images of GBM tumorsphere invasion at day 4 within HA depending on gel degradability (left). Quantification of area changes at day 4 compared to day 0. (right) Statistical significance was analyzed using a two-way ANOVA with Tukey’s multiple comparisons test (**A**, **D**) or an unpaired two-sided Student’s t-test (**C**, **F**). ****p < 0.0001, ***p < 0.001, **p < 0.01.

Given that integrin ligation has been implicated as an important mechanistic step in 3D invasion (*34*), we hypothesized that HA/RGD+ matrices might be able to support faster 3D invasion than HA/RGD-matrices due to the need for matrix secretion in the latter scaffold to support integrin binding. We therefore performed 3D tumorsphere invasion assays to evaluate GBM invasion in RGD- and RGD+ matrices. Indeed, GBM cells invaded the HA/RGD+ gels faster than HA/RGD-gels, achieving 4-fold tumorsphere area changes in 4 days compared to 10 days in HA/RGD-gels (**Fig. 2B**). Moreover, within 4 days, tumorsphere cells in HA/RGD+ gels extended long protrusions into the surrounding matrix whereas TSs in HA/RGD-gels remained highly circumscribed, with few to no discernable protrusions (**Fig. 2C**).

In addition to integrin ligation, proteolytic matrix degradation, particularly due to matrix metalloproteases (MMPs), has been strongly implicated in GBM invasion *in vitro* and *in vivo* (*35*). We therefore reasoned that matrix secretion might depend on MMP-degradability since a non-MMP-degradable matrix might require heightened levels of ECM secretion to facilitate invasion. To test this hypothesis, we compared protein deposition and invasion in 3D HA/RGD-hydrogels crosslinked either with MMP-cleavable (HD/RGD-) or matched non-degradable (ND/RGD-) peptides. Like HA/RGD-matrices, GBM cells in ND/RGD-matrices exhibited significant protein secretion. In contrast, there was substantially less protein secretion from cells in HD/RGD-, with cells in HD/RGD-matrices developing extensive protrusions within 7 days (**Fig. 2D**, fig. S4A). Consistent with our earlier results (**Fig. 1**), GBM cells in stiffer HA/ND/RGD-matrices secreted less protein than cells in soft matrices. Interestingly, this trend reversed in HD/RGD-matrices, with stiffer HD gels supporting slightly greater protein secretion than softer gels. As expected, GBM tumorsphere invasion was higher in HD/RGD-gels than in ND/RGD-gels over 10 days, confirming the importance of MMP degradation in facilitating 3D invasion (**Fig. 2E, F**).

To investigate the combined effects of integrin ligation and MMP degradability on invasion, we tested GBM invasion in HA hydrogels featuring both RGD peptides and MMP-cleavable crosslinkers (fig. S5A). In line with our previous findings, soft HD/RGD+ gels supported less protein secretion than soft ND/RGD+ gels, with the trends reversing for stiff gels (fig. S5B). GBM TSs in both ND/RGD+ and HD/RGD+ gels exhibited protrusions, with longer protrusions and more aggressive invasion in the HD gels (fig. S5C, D).

Collectively, our results show that integrin ligation and MMP degradation both strongly facilitate 3D GBM invasion. When one or both of these mechanisms is initially absent, ECM protein secretion increases, suggesting that protein secretion is a critical adaptive mechanism that renders the microenvironment more hospitable to invasion.

### GBM cultured in HA hydrogels secrete a diverse ensemble of matrisomal proteins whose composition depends on matrix integrin ligation and protease degradability

To identify the proteins produced by GBM cells preceding invasion into 3D RGD-free matrices and the subset of this secretome that includes cell-derived ECM proteins, we performed unbiased mass spectrometric analysis. After cell encapsulation within 3D HA/RGD-gels for defined times, we decellularized hydrogels and harvested the remaining proteins for proteomic analysis (**Fig. 3A**). We used label-free quantification based on spectral counting and calculated the normalized spectral abundance factors (NSAF) to determine relative protein abundances. Comparing our results against the Human Matrisome Database (*36*), we classified our hits into core ECM or ECM-associated proteins.

**Figure 3.**
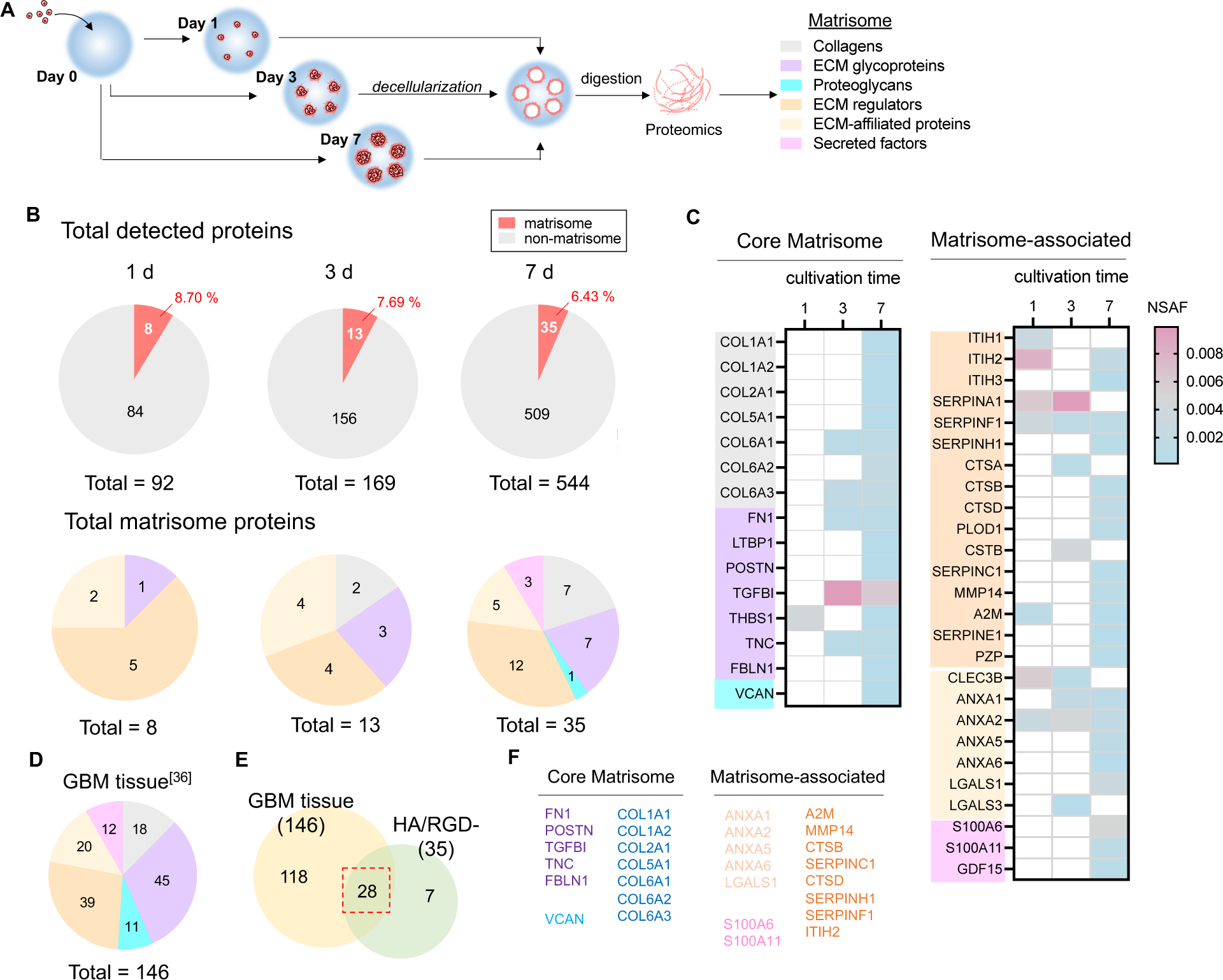
ECM proteins secreted by GBM cells in HA gels are diversified over time and show high similarity to ECM detected from GBM tissue. (**A**) Schematic illustration of the sample preparation process for proteomic analysis. (**B**) Pie charts of the total secreted proteins and total matrisome proteins from the 3D decellularized HA/RGD-gels after 1-, 3- and 7-day GBM cell encapsulation detected by mass spectrometry. (**C**) Heatmaps comparing the matrisomal proteins detected from the decellularized HA/RGD-over time. (**D**) Pie chart showing the compositions of matrisomal proteins from reported GBM tissue proteomics. (**E**) Venn diagram showing the matrisomal proteins in GBM tissue and 3D decellularized HA/RGD-gels. (**F**) Lists of the matrisomal proteins detected from both GBM tissue and 3D decellularized HA/RGD-gels.

To assess the efficiency of our decellularization protocol, we measured DNA content in cell-laden hydrogels at 7 days post-encapsulation in soft and stiff HA/RGD-hydrogels before and after decellularization. Our analysis revealed > 90% reduction in DNA content, confirming the effective removal of cellular contents (fig. S6A, B). Using a colorimetric assay to measure the concentration of secreted peptides, we found we could harvest up to ∼1 mg/mL of the peptide from the decellularized soft HA gels, with a significantly lower concentration (∼0.4 mg/mL) from the stiff gels after decellularization (fig. S6C), consistent with our earlier metabolic labeling measurements (**Fig. 1D**). We also confirmed sample-to-sample reproducibility using three independent biological replicates, which revealed highly overlapping hits in both core matrisome and matrisome-associated proteins (fig. S7).

By comparing proteomic measurements obtained over 1-, 3-, and 7-days post-encapsulation, we found that cells secreted an increasingly diverse ensemble of proteins over time (**Fig. 3B**). Interestingly, while the percentage of secreted proteins that were matrisomal fell with time, the diversity of matrisomal proteins increased. Specifically, 8 out of 92 secreted proteins (8.7%) were matrisomal after 1 day, with that fraction decreasing to 13/169 (7.69%) at 3 days, and 35/544 (6.43%) at 7 days (**Fig. 3B**). Despite the reduction in matrisomal fraction, matrisomal diversity increased with time. At 1 day post-encapsulation, we detected 8 matrisome proteins primarily consisting of ECM regulators (1 glycoprotein, 5 ECM regulators, and 2 ECM-affiliated proteins). By day 3, we identified 13 matrisomal proteins, including 5 core matrisome proteins (2 collagens, and 3 glycoproteins), and 8 ECM-associated proteins (4 ECM regulators, and 4 ECM-affiliated proteins). By 7 days, we detected a total of 35 matrisomal proteins, including 15 core matrisomal proteins (7 collagens, 7 glycoproteins, and 1 proteoglycan) and 20 ECM-associated proteins (12 ECM regulators, 5 ECM-affiliated proteins, and 3 secreted factors). The core matrisomal proteins identified on day 7 contained a variety of integrin-binding ECM proteins, such as collagens, fibronectin, transforming growth factor beta induced protein, and tenascin. These findings support the notion that in the absence of pre-existing integrin-binding motifs, cells secrete integrin-binding proteins in a time-dependent fashion. (**Fig. 3C**).

To investigate the disease relevance of the ECM components we detected in our reductionist system, we compared our findings against published proteomic analyses of human GBM tumors (*37*). To better understand which matrix formulation best matched patient data, we expanded our proteomic analysis to include the RGD-ligating and MMP-degradable/non-degradable gels considered earlier (**Fig. 2**). We found the ECM matrisome composition depended on matrix properties, with the soft HA/RGD-gels at 7 d showing high similarity to brain tissue in terms of total matrisomal content and composition (**Fig. 3B, D**, fig. S8). Specifically, patient tissue and HA/RGD-gels contained 28 common matrisomal proteins (**Fig. 3E, F**). We therefore focused exclusively on soft HA/RGD-gels for subsequent experiments and analyses. Overall, our results show that GBM cells remodel HA matrices over time through ECM secretion, with the greatest remodeling observed for soft, RGD-free, and non-proteolytically degradable matrices.

### Patient transcriptomic analysis identifies upregulation of COL6A3 with increasing glioma grade and in the microvascular proliferation compartment of GBM

We next asked whether the changes we observed in matrix secretion might be reflected in transcriptional data from invasive regions of human GBM tumors. To answer this, we investigated transcriptomic profile changes related to ECM remodeling in patients with GBM using the Rembrandt and Cancer Genome Atlas (TCGA) patient databases. Using the GlioVis data portal (http://gliovis.bioinfo.cnio.es/), we stratified transcriptomic data in terms of tumor grade: non-tumor (NT), low-grade gliomas (LGG), and GBM, enabling us to correlate expression with disease severity.

We first compared differentially expressed genes (DEGs) between NT and GBM tissues using both databases to identify genes upregulated in GBM tissues. Volcano plots of DEGs revealed that compared with NT tissue, approximately 220 (1.11%) of the total 19738 genes in Rembrandt and 771 (6.07%) of the total 12701 genes in TCGA were upregulated in GBM relative to NT tissue (**Fig. 4A, B**). Between these two sets of upregulated genes, 171 genes were enriched in both Rembrandt and TCGA, including 41 matrisome genes (**Fig. 4C**). Gene ontology (GO) analysis of these 171 overlapping upregulated genes revealed close overlap with pathways related to ECM synthesis, assembly, and remodeling (**Fig. 4D**). We repeated the transcriptomic analysis with both databases to identify DEGs between LGG and GBM tissues, which revealed 205 genes commonly enriched in GBM in both databases, of which 47 were matrisomal (fig. S9A, **Fig. 4E**). Of these 205 genes, 34 genes are upregulated in GBM tissue vs. NT, with 17 of these remaining genes being matrisomal (**Fig. 4F**, fig. S9B).

**Figure 4.**
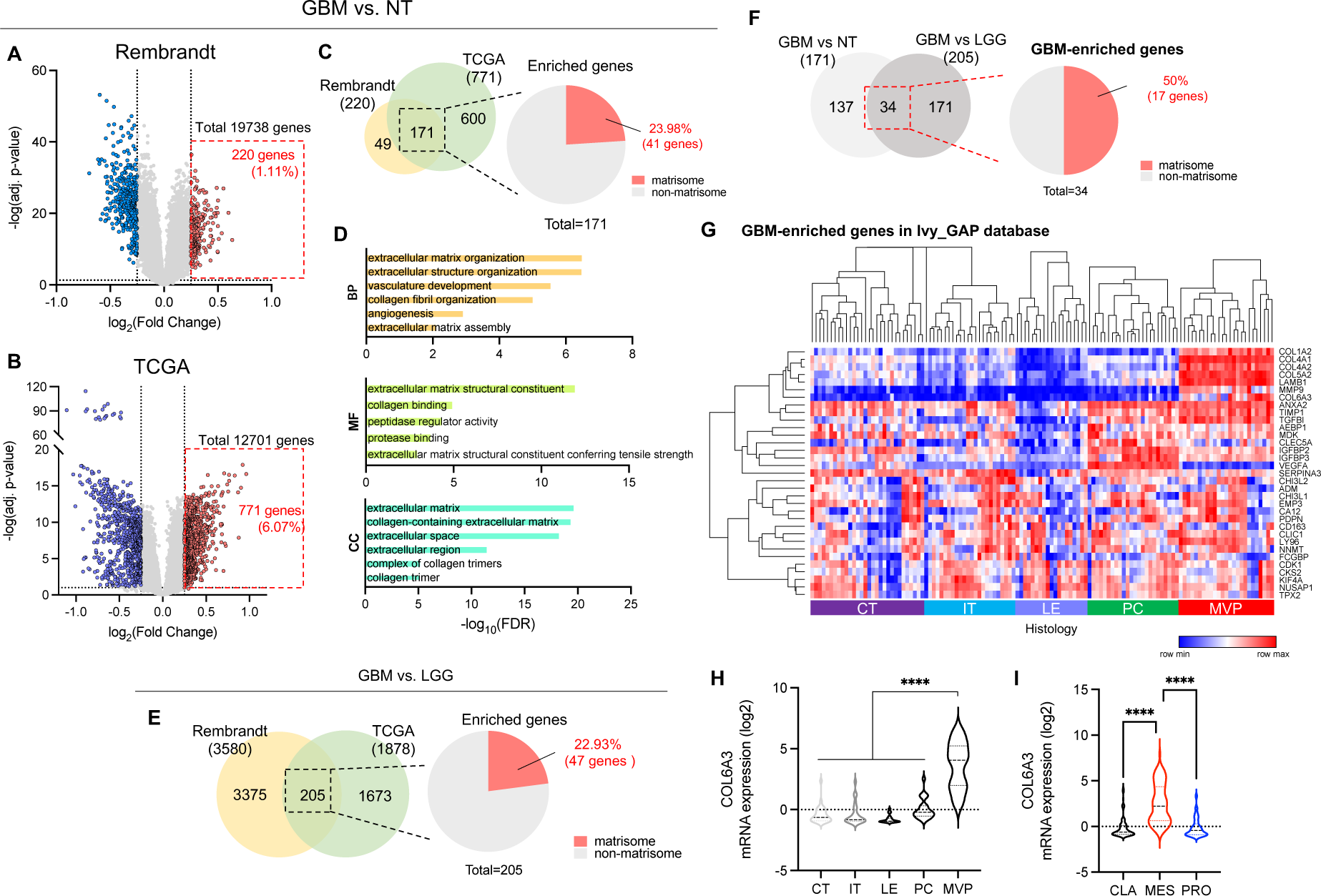
Transcriptomic profiles of GBM patients show significant matrisomal changes during tumor progression. (**A**-**D**) Identification of genes enriched in GBM tissues over non-tumor (NT) tissues using Rembrandt and TCGA databases. Volcano plots from (**A**) Rembrandt and (**B**) TCGA to identify the genes elevated in GBM, compared to NT. (**C**) Venn diagram of genes elevated in GBM compared to NT identified in both Rembrandt and TCGA and pie chart showing the percentage of matrisomal genes in overlapped 171 genes. (**D**) Gene Ontology (GO) analysis of enriched genes from (**C**) (BP: biological process, MF: molecular function, CC: cellular component). (**E**) Venn diagram of genes elevated in GBM compared to low-grade glioma (LGG) identified in both Rembrandt and TCGA and pie chart showing the percentage of matrisomal genes in overlapped 205 genes. (**F**) Venn diagram showing GBM-enriched genes elevated in GBM over both NT and LGG in both Rembrandt and TCGA and pie chart showing the percentage of matrisomal genes in overlapped 34 GBM-enriched genes. (**G**) Heatmap showing region-specific transcriptome profiles of 34 GBM-enriched matrisomal genes. COL6A3 mRNA expression from Ivy_GAP depending on (**H**) histology: cellular tumor (CT), Invasive tumor (IT), leading edge (LE), pseudopalisading cells (PC), and microvascular proliferation (MVP) and (**I**) GBM subtype: classical (CLA), mesenchymal (MES), and proneural (PRO). Elevated genes were identified using a false-discovery rate (FDR) cut-off of < 0.05 and log_2_(fold change) > 0.25 from both enriched gene analyses of GBM vs. NT and GBM vs LGG. Statistical significance was analyzed using one-way ANOVA followed by Tukey’s multiple comparisons test, ****p < 0.001.

The preceding analyses focus on bulk comparisons between different types of tissue, without regard to tumor microanatomy. Thus, we next asked how the expression of matrisomal genes might vary across different tumor microregions, particularly those associated with invasion and chemoresistance. We assessed these microanatomical correlations using the Ivy Glioblastoma Atlas Project (Ivy GAP, http://glioblastoma.alleninstitute.org), which includes transcriptomic data from different regions of patient tumors. We analyzed the regional expression of the 34 genes we previously found to be upregulated in GBM vs NT/LGG tissue and performed hierarchical clustering (**Fig. 4G**). As expected, the expression of these genes varied strongly with histological region. We chose to focus on the region of microvascular proliferation microvascular proliferation (MVP), given the close association of vasculature with GBM infiltration and chemoresistance. In particular, 10 of the genes that were enriched in the MVP region relative to other histological regions were matrisomal, including collagen subunits (COL1A2, COL4A1, COL4A2, COL5A2, and COL6A3), ECM glycoproteins (LAMB1, TGFBI), and ECM regulators (ANXA2, MMP9, and TIMP1). In particular, 10 of the genes that were enriched in the MVP region relative to other histological regions were matrisomal, including collagen subunits (COL1A2, COL4A1, COL4A2, COL5A2, and COL6A3), ECM glycoproteins (LAMB1, TGFBI), and ECM regulators (ANXA2, MMP9, and TIMP1).

To further narrow the list of candidates, we looked for common hits across our transcriptomic and proteomic data. This analysis led us to focus on COL1A2, COL6A3, TGFBI, and ANXA2 as targets strongly enriched in the GBM matrisome at both the transcriptomic and proteomic levels (**Fig. 3F, 4F**, and **4G**). Within this intersection of data sets, COL6A3 is the only matrisomal gene strongly upregulated in the MVP region relative to other regions (**Fig. 4H**, fig. S10). Moreover, COL6A3 was also strongly upregulated in mesenchymal GBMs relative to proneural and classical GBMs, indicating a correlation between COL6A3 expression and tumor aggressiveness (**Fig. 4I**). Based on the prominence of COL6A3 in our *in vitro* studies and our analysis of multiple patient databases, we focused on COL6A3 as a potentially crucial secreted matrisomal protein that primes the brain microenvironment for invasion.

### ECM secretion by GBM cells is highly correlated with invasion associated with bevacizumab resistance

A key finding from our studies above is that COL6A3 is strongly enriched in the region of microvascular proliferation. This zone is commonly associated with tumor angiogenesis, which is a hallmark of GBM and involves extensive remodeling of endogenous ECM to form new blood vessels. These neovessels are often disorganized and leaky, causing hypoxia and a shift towards a mesenchymal phenotype with increased tissue invasion (*38*). Transition to a mesenchymal phenotype is an important feature of bevacizumab treatment and resistance (*39*), where tumor cells that are deprived of the ability to induce angiogenesis invade tissue and home towards vasculature. This process is mediated in part by the transcription factor ZEB1, a known driver of the mesenchymal phenotype (*25*). We therefore hypothesized that ECM deposition in general and collagen VI deposition in particular might functionally contribute to the enhanced invasion associated with resistance to bevacizumab and potentially other anti-angiogenic agents in GBM.

To investigate this possibility, we performed comparative studies between our previously described U87 GBM cells conditioned to be bevacizumab-resistant (Bev^R^) and their matched bevacizumab-sensitive counterparts (Bev^S^) (*25*). Notably, Bev^R^ cells recapitulate the invasive phenotype seen in human GBMs following bevacizumab resistance (*40*). While both Bev^S^ and Bev^R^ cells developed micro-protrusions within the HA/RGD-matrices after 7 days of culture, Bev^R^ cells formed more micro-protrusions than Bev^S^ cells, as observed by differential interference contrast imaging (fig. S11A, B). Moreover, 3D tumorsphere invasion assays revealed that Bev^R^ cells invaded HA gels to 1.2-fold higher areas than Bev^S^ cells after 4 days (fig. S11C). Because the greater protrusive activity of the Bev^R^ cells suggested a more active interaction with the HA matrix, we hypothesized that Bev^R^ cells might secrete more ECM to support greater matrix engagement and invasion. Consistent with this hypothesis, we observed that Bev^R^ cells secreted more ECM than Bev^S^ cells by metabolic labeling (**Fig. 5A**). In addition, we compared the changes in matrix stiffness after 7 days of Bev^S^/Bev^R^ cell encapsulation (**Fig. 5B**), which revealed an increase in matrix Young’s modulus for Bev^R^ cell-laden gels.

**Figure 5.**
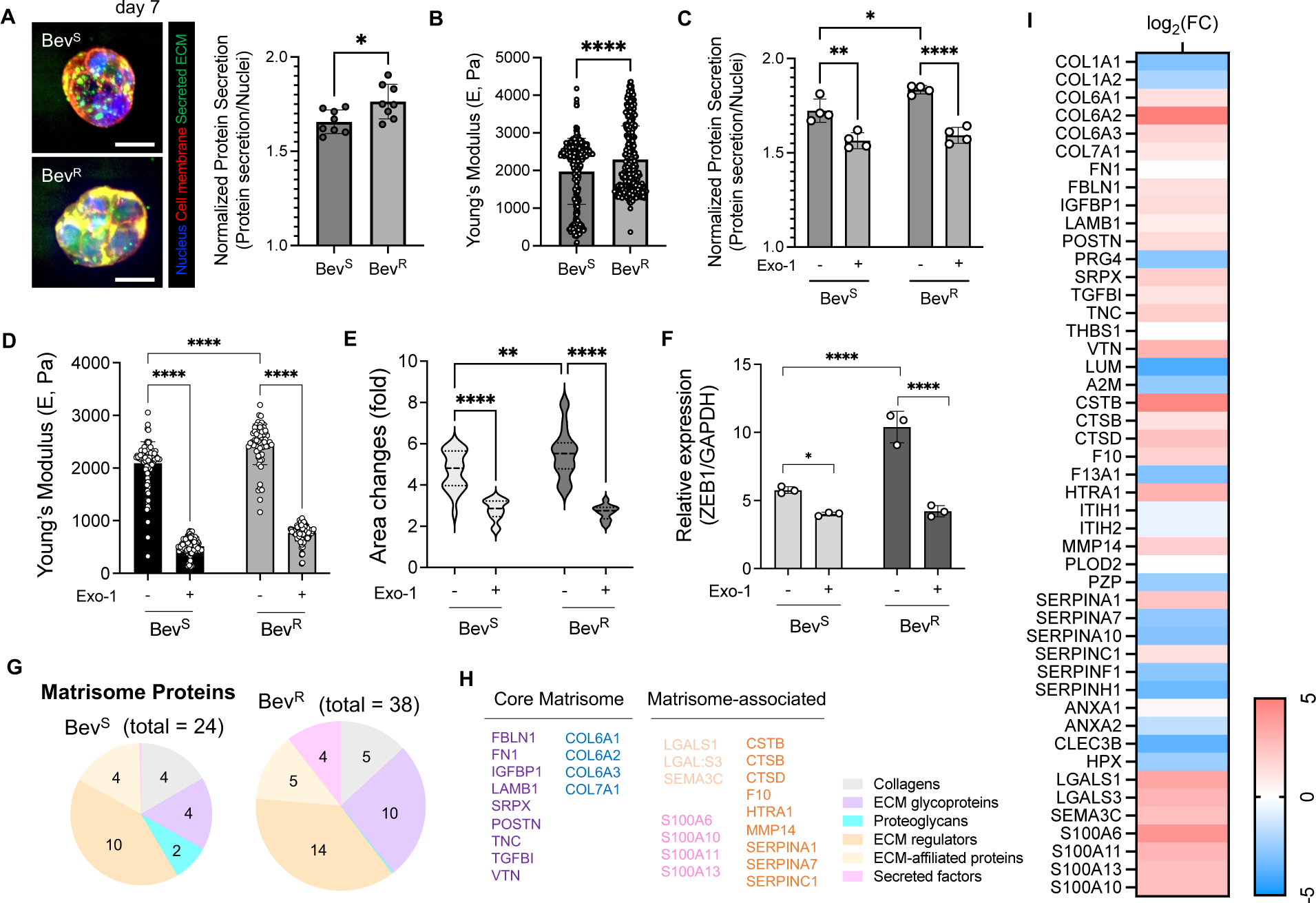
ECM secretion and matrix stiffness are elevated in bevacizumab-resistant GBM cells in HA gels. (**A**) Representative fluorescence images (left) and quantification (right) of secreted ECM by Bev_S_ and Bev_R_ cells within HA/RGD-gels at day 7. Scale bar: 30 μm. (n = 8) (**B**) Comparison of Young’s modulus of Bev^S^ and Bev^R^ cell-laden gels at day 7. (**C**) Quantification of secreted ECM by Bev^S^ and Bev^R^ cells within 3D HA/RGD-gels when treated with Exo-1 (inhibitor of exocytosis and vesicular trafficking, 100 nM). (n = 4) (**D**) Comparison of Young’s modulus of Bev^S^ and Bev^R^ cell-laden HA/RGD-with Exo-1 treatment at day 7. (**E**) Quantification of 3D sphere invasion assay for Bev^S^ and Bev^R^ at day 4, in response to Exo-1 treatment. (n = 20 to 21) (**F**) ZEB1 gene expression in Bev^S^ and Bev^R^ cells within 3D HA/RGD-when treated with Exo-1. (n = 3) (**G**) Pie charts of matrisomal proteins identified from decellularized 3D HA/RGD-after Bev^S^ and Bev^R^ cell encapsulation. (**H**) List of matrisomal proteins highly detected from Bev^R^ cell-laden HA/RGD-gels, compared to Bev^S^ cell-laden gels. (**I**) Average log_2_ fold change (FC) difference of matrisomal proteins secreted by Bev^R^ cells over Bev^S^ cells within 3D HA/RGD-by proteomics. Statistical significance was analyzed using an unpaired two-sided Student’s t-test (**B**, **C**) or a two-way ANOVA with Tukey’s multiple comparisons test (**C**∼**F**). ****p < 0.0001, **p < 0.01, *p < 0.05.

The preceding data establishes a correlation between ECM protein secretion and HA stiffening. To explore potential causal relationships, we repeated these experiments under inhibition of protein secretion with Exo-1, which limits vesicular trafficking between the endoplasmic reticulum and Golgi, consequently reducing ECM secretion (*41*). As expected, Exo-1 treatment noticeably reduced ECM secretion in both Bev^S^ and Bev^R^ cells (Fig. S12A, **Fig. 5C**). For both Exo-1-treated Bev^R^ and Bev^S^ cell-laden hydrogels, AFM revealed dramatic reductions in Young’s modulus, and spheroid invasion assays showed markedly reduced invasion (Fig. S12B, **Fig. 5D, E**). ZEB1 expression also fell to similar levels in both Bev^S^ and Bev^R^ cells in response to Exo-1 treatment (**Fig. 5F**). Collectively, these findings indicate that ECM secretion underlies altered tumor stiffness and invasion associated with bevacizumab resistance.

We next performed proteomic analysis on the Bev^S^ and Bev^R^ secretome to identify potential mediators of increased invasion. Mass spectrometry revealed 38 matrisomal proteins secreted by Bev^R^ GBM cells including 15 core proteins (5 collagens, 10 glycoproteins) and 23 ECM-associated proteins (14 ECM regulators, 5 ECM-affiliated proteins, and 4 secreted factors), while 24 matrisomal proteins were secreted by Bev^S^ cells, with 10 core proteins (4 collagens, 4 glycoproteins, and 2 proteoglycans) and 14 ECM-associated proteins (10 ECM regulators and 4 ECM-affiliated proteins) (**Fig. 5G**). Strikingly, COL6A1, COL6A2, and COL6A3 subunits were among the most upregulated ECM components in Bev^R^ cells relative to Bev^S^ cells (**Fig. 5H, I**). These findings raise the intriguing possibility that matrix remodeling driven by matrix protein secretion, especially of collagen VI, promotes GBM invasion associated with bevacizumab resistance.

### Secreted collagen VI colocalizes with integrins and triggers β-catenin signaling and ZEB1 expression

Next, we more deeply investigated the possibility that GBM cells secrete collagen VI to promote invasion as part of the bevacizumab-resistance phenotype. Immunostaining revealed rich collagen VI density within secreted and metabolically labeled ECM, with much greater co-localization in Bev^R^ cells than in Bev^S^ cells (**Fig. 6A**). qPCR of HA-encapsulated single cells confirmed that Bev^R^ cells express ∼3-fold higher collagen VI than Bev^S^ cells (**Fig. 6B**).

**Figure 6.**
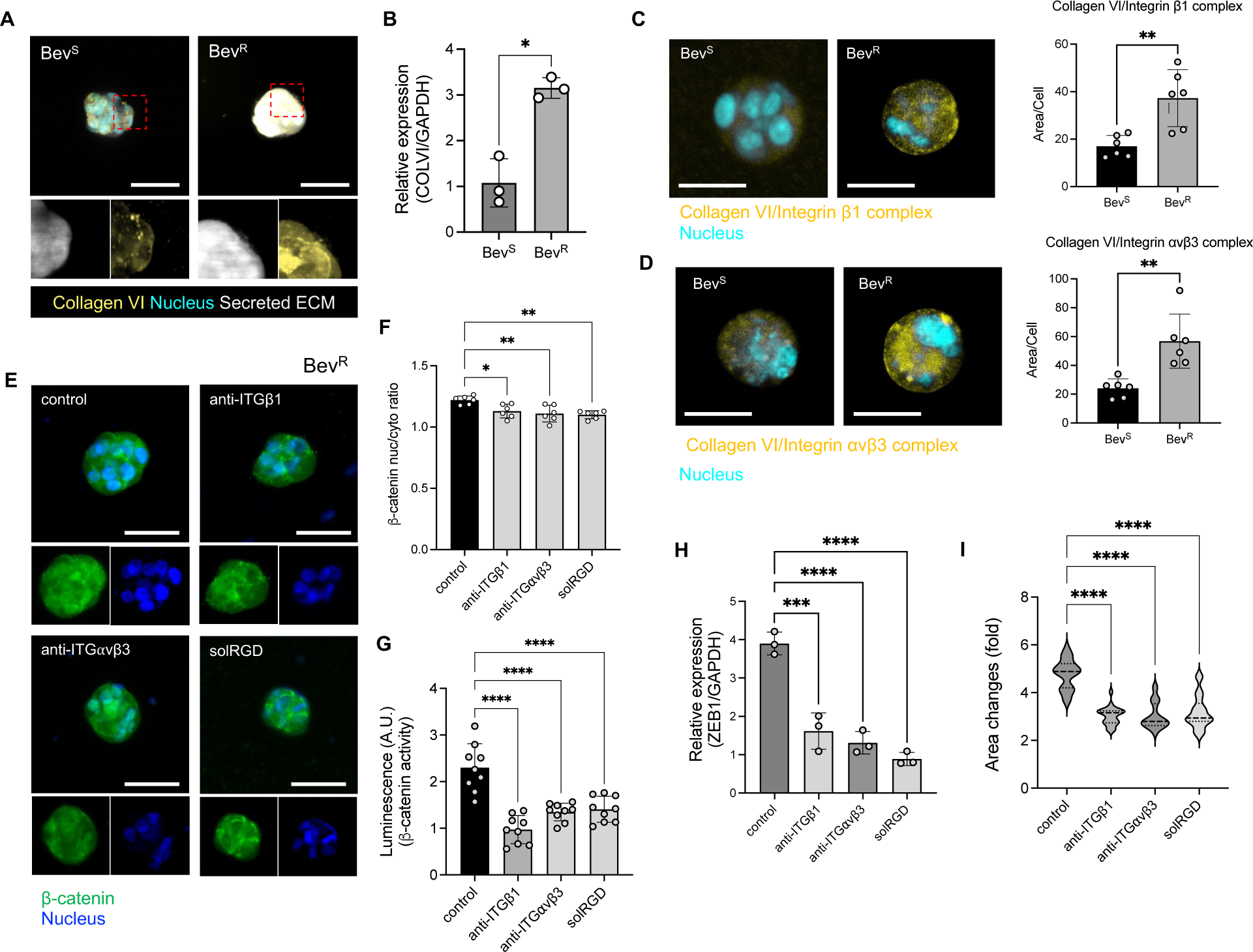
Interactions between secreted collagen VI and integrins regulate p-catenin signaling and ZEB1 expression. (**A**) Representative fluorescent images of secreted ECM and collagen VI from Bev_S_ and Bev_R_ cells within 3D HA/RGD-gels at day 7. Scale bar: 50 pm. Inset: magnified images of the red dotted box. (**B**) Relative mRNA levels of collagen VI in Bev^S^ and Bev^R^ cells within 3D HA/RGD-gels. (n = 3) (**C**, **D**) Proximity ligation assay (PLA) to investigate secreted collagen VI-integrins engagement. (**C**) Representative images (left) and quantification (right) of collagen VI interactions with integrin pl in Bev_S_ and Bev_R_ cells within 3D HA/RGD-gels. Scale bar: 50 pm. (n = 6) (**D**) Representative images (left) and quantification (right) of collagen VI interactions with integrin avp3 in Bev_S_ and Bev_R_ cells within 3D HA/RGD-gels. Scale bar: 50 pm. (n = 6) (**E**) Representative fluorescent images of p-catenin in Bev^R^ cells in HA/RGD-gels after 7 days of no treatment (control), or treatment with monoclonal integrin pi antibodies (anti-ITGpi), integrin avp3 antibodies (anti-ITG avp3), or soluble RGD (solRGD). Scale bar: 50 pm. (**F**) Quantification of p-catenin nuclear localization in Bev^R^ cells in response to inhibition of integrin binding within HA/RGD-gels. (**G**) Luciferase assay to assess p-catenin activity in Bev^R^ cells in response to inhibition of integrin binding within HA/RGD-gels. (n = 9) (**H**) ZEB1 gene expression (n = 3) in Bev^R^ cells and (**I**) quantification of 3D sphere invasion assay of Bev^R^ cells with the perturbation of integrin binding within HA/RGD-gels. (n = 21 to 24) Statistical significance was analyzed using an unpaired two-sided Student’s t-test (**B**∼**D**) or a one-way ANOVA with Tukey’s multiple comparisons test (**F**∼**I**). ****p < 0.0001, ***p < 0.001, **p < 0.01, *p < 0.05.

Based on the results, we then investigated whether the GBM-secreted collagen VI interacts with its cognate integrin receptors. Since several cell surface receptors including the α1β1, α2β1, α3β1, α10β1, and αvβ3 integrins are known to be capable of binding to collagen VI (*42*), we performed proximity ligation assays (PLA) to visualize collagen VI interactions with integrin β1 and integrin αvβ3 in Bev^S^ and Bev^R^ cells. Higher collagen VI PLA colocalization was observed in Bev^R^ cells than in Bev^S^ cells for both integrin β1 and integrin αvβ3 (**Fig. 6C, D**).

Next, we examined whether interactions between secreted collagen VI and its cognate integrins might transduce adhesion-dependent signals key to invasion. Wnt/β-catenin signaling represents a particularly important signaling system given its upregulation in aggressive GBMs, its association with integrin-dependent mechanotransduction (*43, 44*), and its established relevance to the bevacizumab-induced mesenchymal transition in GBM (*45–47*). We therefore hypothesized that secreted collagen VI acts through its cognate integrin receptors to trigger β-catenin signaling to drive invasion.

To address this possibility, we examined β-catenin nuclear localization in both Bev^S^ and Bev^R^ cells. The nuclear/cytoplasmic intensity ratio of β-catenin in Bev^S^ and Bev^R^ cells was measured at 7 days post-encapsulation and found to be higher in Bev^R^ cells than in Bev^S^ cells (Fig. S13), which implies amplification of β-catenin signaling in Bev^R^ cells. To determine whether this heightened β-catenin nuclear localization depended on integrin binding, we repeated our studies in the presence of inhibitory monoclonal integrin β1 antibodies (anti-β1) and the cyclic RGD pentapeptide Cilengitide (anti-αvβ3) to selectively block interactions of each integrin subunit with secreted collagens, with soluble RGD peptides (sol RGD) as an additional and broader competitive inhibitor for comparison. The nuclear/cytoplasmic intensity ratio of β-catenin was significantly reduced across all treatment groups relative to controls (**Fig. 6E, F**). To test the effects of integrin binding on β-catenin-dependent transcription, we performed these inhibition studies in cells expressing a β-catenin luciferase reporter construct (**Fig. 6G**). Relative to the untreated group, we observed significantly less bioluminescence following treatment with all integrin inhibitors, indicating that β-catenin nuclear localization and transcriptional function are driven by integrin binding to secreted ECM components.

To determine whether integrin binding also causes the expression of mesenchymal markers associated with invasion, we measured ZEB1 expression with and without integrin inhibition. Inhibition of integrin binding markedly reduced ZEB1 expression (**Fig. 6H**), which was accompanied by a strong reduction in 3D invasion by spheroid assay (**Fig. 6I**). This upregulation of ZEB1 was strongly dependent on β-catenin signaling, because inhibition of the β-catenin-TCF/LEF interaction (iCRT14) reduced 3D invasion, whereas suppression of β -catenin destruction (via CHIR) increased 3D invasion (fig. S14).

These studies establish that secreted ECM acts through collagen VI-binding integrins to stimulate β-catenin signaling and ZEB1 expression to promote invasion associated with bevacizumab resistance.

### Secreted collagen VI is involved in matrix stiffening and mechanotransductive signaling that promotes GBM invasion associated with bevacizumab resistance

Collagen VI is classically considered a microfibril-forming collagen that can bind different ECM components and ECM-associated macromolecules, including other types of collagens, fibronectin, and HA (*48–51*). In this way, collagen VI is thought to contribute to the organization of 3D tissue architecture in various tissues as a ‘matrix crosslinker.’ This reported role led us to speculate that secreted GBM cell-secreted collagen VI might mechanically stiffen the surrounding ECM and transduce mechanotransductive signals that contribute to GBM invasion associated with bevacizumab resistance (**Fig. 7A**).

**Figure 7.**
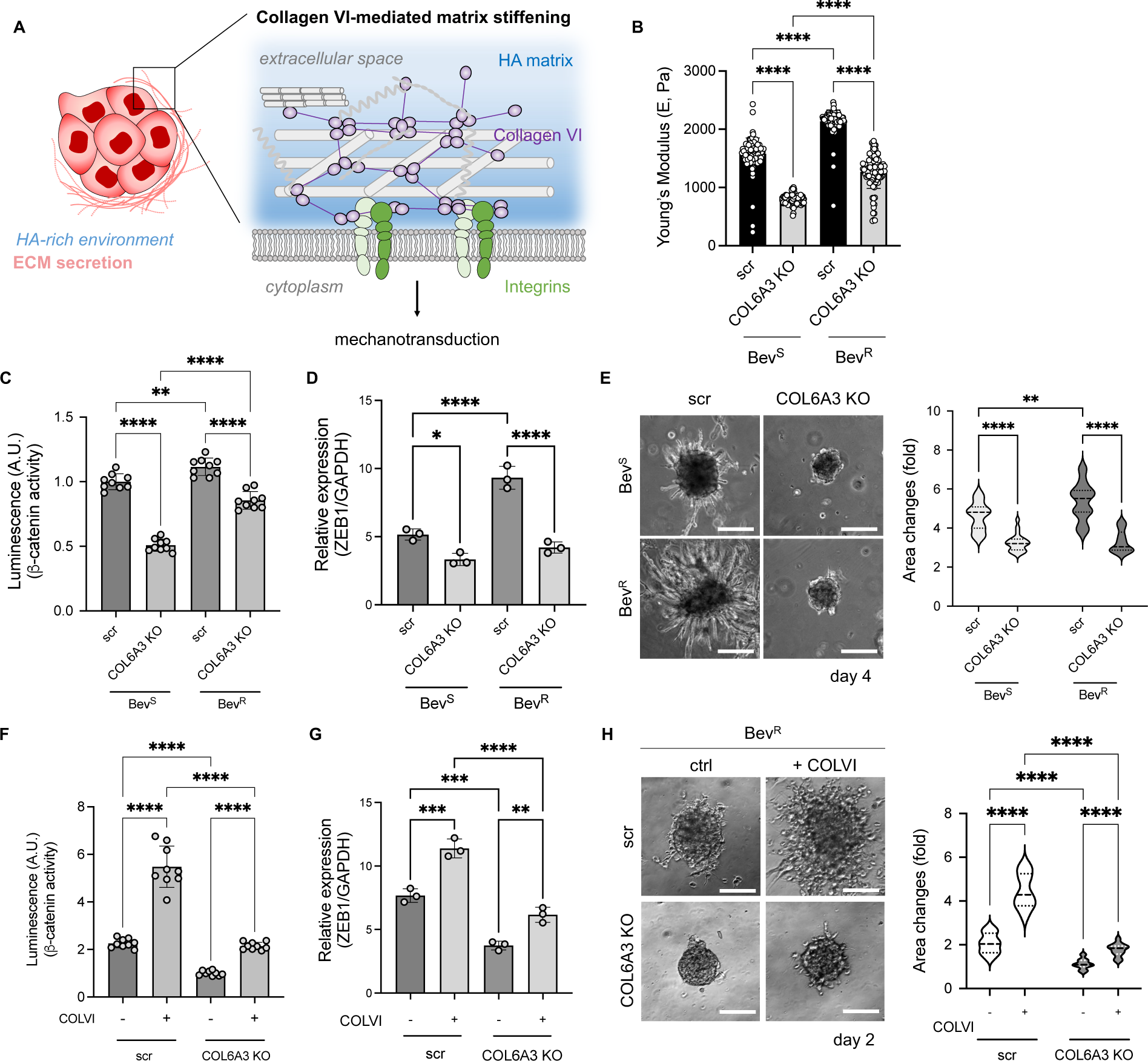
Collagen VI secreted by GBM cells is involved in matrix stiffening and mechanotransductive signaling, regulating bevacizumab resistance-associated invasion. (**A**) Schematic illustration of the hypothetical mechanism of matrix stiffening and mechanotransduction driven by collagen VI secreted by GBM cells. (**B**) Comparison of Young’s modulus of gels encapsulating non-targeting and COL6A3 knockout (KO) Bev^S^ and Bev^R^ cells. (**C**) Luciferase assay for assessing p-catenin activity for nontargeting and COL6A3 KO Bev_S_ and Bev^R^ cells after encapsulating within HA/RGD-gels for 7 days. (n = 9) (**D**) ZEB1 gene expression of nontargeting and COL6A3 KO Bev^S^ and Bev^R^ cells after culturing in HA/RGD-gels for 7 days. (**E**) Representative images (left) and quantification (right) of 3D invasion of nontargeting Bev^S^ and Bev^R^ tumorspheres compared to COL6A3 KO tumorspheres at day 4. Scale bar: 100 pm. (**F**) Luciferase assay for assessing p-catenin activity for nontargeting and COL6A3 KO Bev^R^ cells with or without human collagen VI treatment within HA/RGD-gels for 7 days. (**G**) ZEB1 gene expression of nontargeting and COL6A3 KO Bev^R^ cells with or without human collagen VI treatment cultivated in HA/RGD-gels for 7 days. (**H**) Representative images (left) and quantification (right) of 3D invasion of nontargeting and COL6A3 KO Bev^R^ tumorspheres depending on human collagen VI treatment at day 2. Scale bar: 100 um. Statistical significance was analyzed using a two-way ANOVA with Tukey’s multiple comparisons test. ****p < 0.001, ***p < 0.005, **p < 0.01, *p < 0.05.

Thus, we asked whether the HA matrix stiffening we observed is dependent on cell-secreted collagen VI. We performed CRISPR-based gene editing to create COL6A3 knockout (KO) GBM cells for both Bev^S^ and Bev^R^ cells (fig. S15). AFM revealed that both COL6A3 KO Bev^S^ and Bev^R^ cells-laden gels were significantly softer than their corresponding controls (∼0.5-fold and ∼0.6-fold, respectively) (**Fig. 7B**). Nonetheless, COL6A3 KO Bev^R^-laden HA gels were still stiffer than COL6A3 KO Bev^S^-laden HA gels, implying that the greater overall matrix secretion observed with Bev^R^ cells (fig. S16) may contribute to increases in matrix stiffness.

We therefore asked whether the reductions in stiffness associated with COL6A3 KO translated into reduced Wnt/β-catenin signaling. Using the β-catenin bioluminescence reporter described earlier, we observed significant decreases in β-catenin-dependent transcription in both Bev^S^ and Bev^R^ cells when the COL6A3 gene was deleted (**Fig. 7C**). Consistent with the differential COL6A3 KO-dependent effect on stiffness between Bev^R^ and Bev^S^ cells, Bev^R^ cells showed higher β-catenin-dependent transcriptional activity than Bev^S^ cells. Moreover, COL6A3 KO significantly decreased mRNA expression level of ZEB1 and 3D invasion in both Bev^S^ and Bev^R^ cells relative to non-targeting controls (**Fig. 7D, E**), with little difference in these metrics between COL6A3 KO Bev^S^ and Bev^R^ cells.

To confirm the COL6A3-specificity of these effects, we performed complementation studies in which we added exogenous (purified) collagen VI to COL6A3 KO Bev^R^ cells. Collagen VI supplementation partly rescued β-catenin activity and ZEB1 expression in Bev^R^ cells to similar levels seen in non-supplemented, non-targeting controls (**Fig. 7F, G**). A similar trend was observed in the 3D invasion assay (**Fig. 7H**). These results demonstrate that collagen VI directly contributes to matrix stiffening and subsequent mechanotransduction relevant to the mesenchymal phenotype and invasion associated with bevacizumab resistance.

### Hypoxia amplifies collagen VI-dependent mechanotransduction, ZEB1 expression, and invasion

The GBM microenvironment is characterized by hypoxia due to rapid proliferation, vascular insufficiency, and other factors (*52, 53*). This hypoxia can in turn promote a mesenchymal shift with increased migration and invasion, as seen in the adaptive-evasive response in bevacizumab resistance. Hypoxia can also preserve and promote the GSC population, setting the stage for therapeutic resistance and recurrence (*54*). Given our results demonstrating that collagen VI contributes to bevacizumab resistance-associated invasion, we asked whether this effect might be amplified in the setting of hypoxia.

We began by repeating proteomic analysis of secreted ECM components by Bev^S^ and Bev^R^ GBM cells under normoxic (20 % O_2_) and hypoxic (1 % O_2_) conditions (fig. S17A). Interestingly, hypoxia induced very distinct core matrisomal profiles in Bev^S^ and Bev^R^ cells. For instance, some core matrisomal proteins such as COL6A1, COL6A3, TGFBI, TNC, and VTN were enriched in hypoxic Bev^R^ cells relative to their normoxic counterparts. In contrast, matrisome-associated proteins were less abundantly detected under hypoxic conditions in both Bev^S^ and Bev^R^ cells.

To determine whether collagen VI expression changes with hypoxia, we immunostained for collagen VI in both Bev^S^ and Bev^R^ cells under normoxic and hypoxic conditions (**Fig. 8A**). Hypoxia produced more intense collagen VI intensity in both Bev^S^ and Bev^R^ cells, with greater collagen VI secretion in Bev^R^ cells under both conditions. qPCR confirmed similar trends in collagen VI transcript levels in Bev^S^ vs. Bev^R^ cells and hypoxia vs. normoxia (**Fig. 8B**). PLA revealed enhanced collagen VI-integrin interactions under hypoxic conditions (fig. S17B, C). Correspondingly, hypoxia increased β-catenin nuclear translocation in both Bev^S^ and Bev^R^ cells (**Fig. 8C**) as well as ZEB1 expression and invasion (**Fig. 8D, E**), consistent with a hypoxia-induced mesenchymal transition. All three effects could be reduced by antibody-based blockade of integrin of β1 and αvβ3 (Fig. S18), indicating that hypoxia-induced collagen VI acts through specific integrins to promote a mesenchymal phenotype characterized by enhanced invasion.

**Figure 8.**
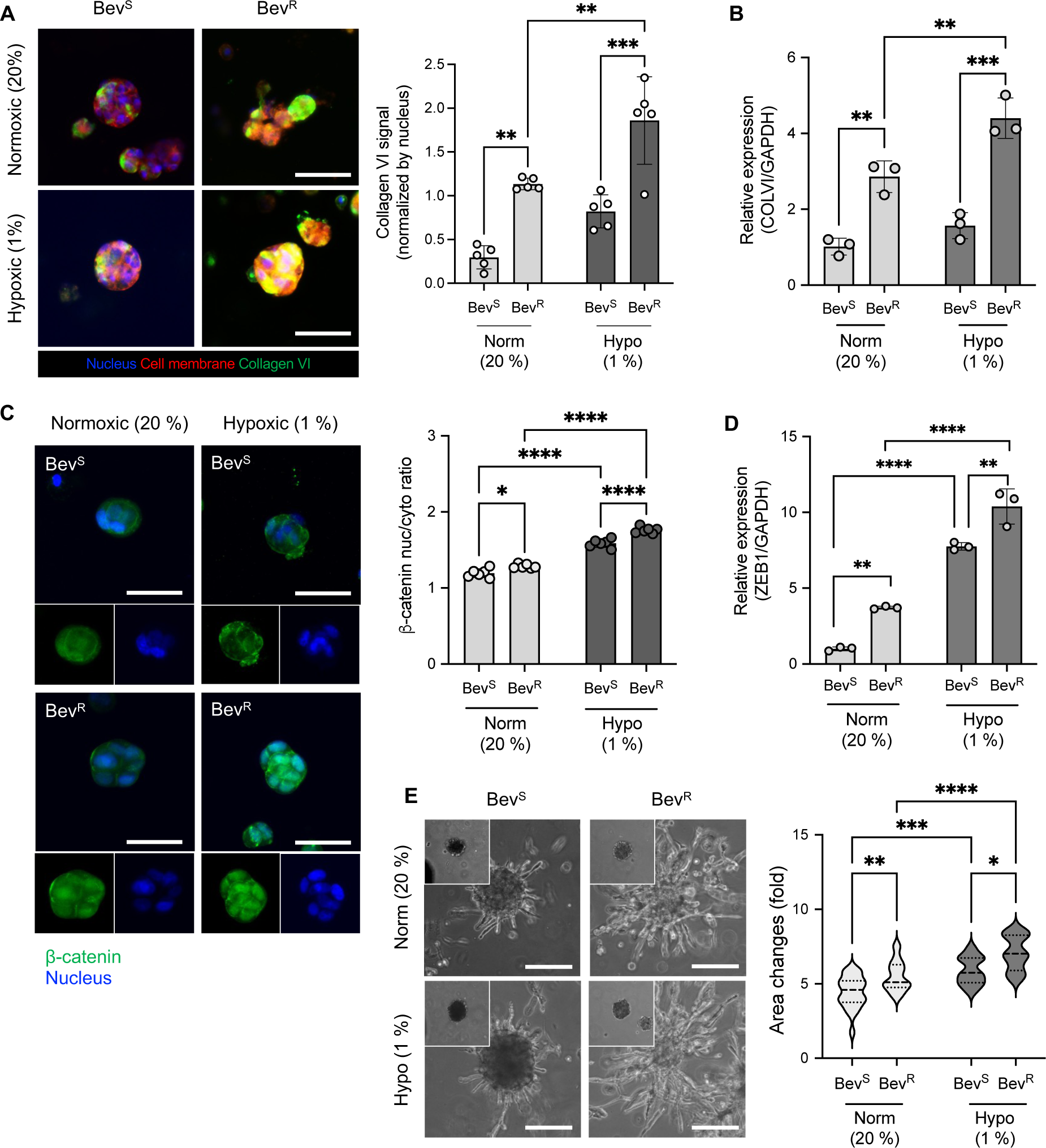
Secreted collagen VI-triggered mechanotransduction is highly enhanced in hypoxic environments, promoting invasion. (**A**) Representative fluorescent images (left) and quantification (right) of collagen VI in Bev^S^ and Bev^R^ cells within 3D HA/RGD-gels under normoxic (20%) and hypoxic (1%) conditions. Scale bar: 50 pm. (**B**) Collagen VI gene expression in Bev^S^ and Bev^R^ cells within 3D HA/RGD-gels under normoxic (20%) and hypoxic (1%) conditions. (**C**) Representative fluorescence images (left) and quantification of p-catenin nuclear translocation (right) of p-catenin in Bev_R_ cells within HA/RGD-gels under normoxic (20%) and hypoxic (1%) conditions after 7 days. Scale bar: 50 pm. (**D**) ZEB1 gene expression in Bev^S^ and Bev^R^ cells within 3D HA/RGD-gels under normoxic (20%) and hypoxic (1%) conditions. (**E**) Representative images (left) and quantification (right) for 3D invasion of Bev^S^ and Bev^R^ tumorspheres under normoxic (20%) and hypoxic (1%) conditions at day 4. Insets: tumorspheres at day 0. Scale bar: 100 um. Statistical significance was analyzed using a two-way ANOVA with Tukey’s multiple comparisons test. ****p < 0.001, ***p < 0.005, **p < 0.01, *p < 0.05.

Finally, to test relationships between collagen VI production and the mesenchymal phenotype in a more patient-proximal model, we conducted a comparative study on patient-derived GSCs classified as proneural (GSC262) and mesenchymal (GSC20). We encapsulated both cells within HA matrices to test the effect of GBM subtypes on ECM remodeling (fig. S19). GSC20 cells produced elongated protrusions into the surrounding matrix, while GSC262 cells grew in a more circumscribed fashion, without appreciable protrusive engagement of the matrix (fig. S19A). GSC20 cells exhibited much greater overall matrix secretion than GSC262 cells (fig. S19B), as well as greater COLVI and ZEB1 expression (Fig. S19C, D).

## Discussion

Our study explores how GBM cells secrete matrix proteins to convert the surrounding microenvironment from normal ECM into an ECM highly supportive of GBM invasion. Using a combination of metabolic labeling and mass spectrometry, we found that the extent and composition of secreted matrisome depend on matrix stiffness, integrin ligation, and proteolytic degradability, with the greatest matrix secretion seen in the matrices that are initially least supportive to invasion. By correlating our proteomic data with transcriptomic data from patient GBMs, we identify collagen VI as a potential mediator of invasion. We then go on to show that secreted collagen VI both stiffens the matrix and ligates specific integrins, promoting mechanotransductive signaling that stimulates β-catenin signaling, ZEB1 expression, and invasion. These effects are strongest in GBM cells conditioned to be bevacizumab-resistant and/or cultured under hypoxic conditions (**Fig. 9**).

**Figure 9.**
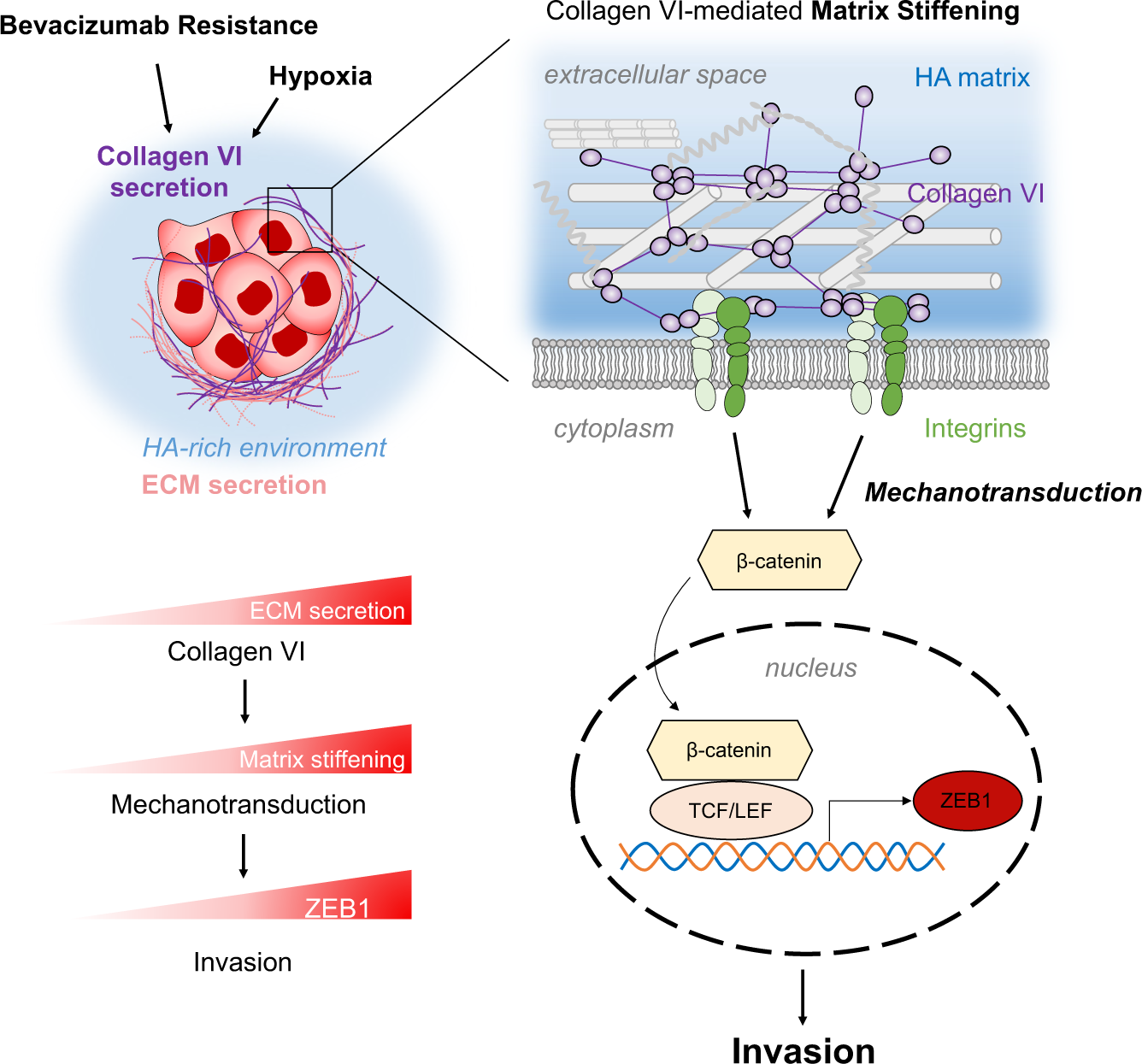
Schematic illustration of the proposed mechanism. Matrix stiffening driven by collagen VI secreted by GBM cells regulates ZEB1 through β-catenin signaling, contributing to the invasion associated with bevacizumab resistance and hypoxia in GBM.

Perhaps the most important and unexpected finding from our study is the importance of collagen VI in priming the microenvironment for invasion. Collagen VI has been highlighted as both a biomarker and functional driver of various disease states including muscular dystrophy, fibrosis, and multiple cancer types (*55, 56*). In GBM specifically, at least one study has also found a causative role for collagen VI in driving tumor angiogenesis and progression in a mouse xenograft model (*10*). Our findings support and build upon that work by implicating collagen VI in the remodeling of the matrix to facilitate invasion, which involves matrix stiffening and mechanotransductive signaling through integrins to promote mesenchymal gene expression and invasion. Importantly, collagen VI secretion precedes invasion and is most enhanced in matrices that are least amenable to invasion (non-RGD-conjugated and non-proteolytically degradable), strongly suggesting that GBM cells secrete collagen VI as an adaptive mechanism to prime the microenvironment for invasion. HA matrices were an important enabling tool in our work, both because this material allows modular tuning of mechanics, integrin ligation, and enzymatic degradability and because HA is the most abundant matrix component in the brain. The latter point is especially significant, because collagen VI is known to bind HA as part of its function as a matrix crosslinker within tissue (*48*), with HA/aggrecan complexes proposed to play a direct role in organizing the assembly and architecture of collagen VI microfibrillar networks (*57*).

Our study also raises the question of what structural role collagen VI plays in promoting invasion. Collagen VI is typically classified as microfibril-forming collagen, where it assembles from triple-helical precursors to form beaded microfilaments. The collagen VI α3 chain is essential for this assembly and deposition process (*58*), which aligns well with our finding that collagen VI α3 is enriched in the GBM secretome and functionally important to invasion. The collagen VI α3 chain contains a rich range of functional domains, including a fibronectin-type-III domain and a Kunitz-like domain. Collagen VI α3 can also be proteolyzed to yield the cleavage product endotrophin (ETP), which can itself stimulate cell migration and metastasis (*59*) by enhancing transforming growth factor β (TGF-β) signaling to promote mesenchymal transition. While our studies make clear that collagen VI can promote invasion through integrin-based mechanotransduction, it will be valuable in future studies to elucidate contributions via ETP and other degradation products.

Finally, our study helps form conceptual links between ECM remodeling, mesenchymal transition, and bevacizumab-resistant invasion in GBM. In previous work, we demonstrated an association between GBM bevacizumab resistance and a ZEB1-mediated mesenchymal transition in GBM, ultimately driving perivascular invasion (*25*). Consistent with our earlier study, our transcriptomic analysis revealed strong upregulation of pathways related to angiogenesis and associated ECM remodeling, which are predictive of the clinical response to bevacizumab (*60*). Our study goes beyond these correlations and directly implicates collagen VI secretion in ECM remodeling key to the invasive phenotype associated with bevacizumab resistance. This relationship raises the exciting possibility that collagen VI and its cognate integrins could potentially be therapeutically targeted to limit invasion associated with bevacizumab resistance and thus improve clinical outcomes.

## Materials and Methods

### HA hydrogel synthesis

Methacrylated HA (Me-HA) was synthesized as described previously (*30*). Briefly, methacrylic anhydride (Sigma-Aldrich) was used to functionalize sodium hyaluronate (Lifecore Biomedical, HA60K, molecular weight: 66 kDa–99 kDa) with methacrylate groups. To add integrin-adhesive functionality, Me-HA was conjugated via Michael Addition with the integrin-binding RGD peptide Ac-GCGYGRGDSPG-NH2 (Anaspec) at a final concentration in the gel of 0.5 mmol/L. Finally, 1.5 wt/wt % Me-HA was crosslinked in phenol-free DMEM (Gibco) with bifunctional thiol dithiothreitol (DTT, Thermo Fisher Scientific) or a peptide crosslinker that was either broadly protease-degradable (KKCGGPQGIWGQGCKK, Genscript) or non-degradable (KKCGGDQGIAGFGCKK, Genscript). The value of thiol group : HA monomer ratio was selected to yield a Young’s modulus of ∼0.25 kPa and ∼2.5 kPa for soft and stiff conditions, respectively. The mixture was incubated for 1 h to be crosslinked at 37 °C, 5% CO_2,_ and immersed in a cell culture medium.

### Cell culture

U87 MG human glioblastoma cells were obtained from the University of California, Berkeley Tissue Culture Facility, which obtains its cultures from the American Type Culture Collection (ATCC). Cells were cultured in fully supplemented high-glucose DMEM (Invitrogen) supplemented with 10% fetal bovine serum (Corning), 1× MEM non-essential amino acids solution (Gibco), 1mM sodium pyruvate (Gibco), and 1% penicillin-streptomycin (Gibco). Cells were maintained at 37 °C, 5% CO2, and dissociated using 0.25% trypsin-EDTA (Thermo Fisher Scientific) for passaging every 3∼4 days. Bevacizumab-sensitive/resistant U87 cells were cultured and passaged identically to wild-type U87 cells (*25*). Cells were screened for mycoplasma and authenticated every 6 months by Short Tandem Repeat (STR) analysis at the UC Berkeley Cell Culture Facility.

For 3D encapsulation of individual cells within HA gels, dissociated cells were resuspended at the desired cell density in phenol-free DMEM and added to the HA mixture before cross-linking. Cell-laden HA gels were inspected via phase contrast imaging (Eclipse TE2000-E, Nikon) and images of encapsulated cells were quantified to calculate the shape index within HA gels (4ν x area/perimeter^2^) using ImageJ. For visualization of secreted proteins, cell-laden hydrogels were cultured in AHA growth media, which consists of DMEM, high glucose, no glutamine, no methionine, no cystine (Gibco) supplemented with 0.2 mM AHA (Click chemistry tools), 4 mM L-glutamine (Gibco), 0.201 mM cystine (Sigma-Aldrich), 10% fetal bovine serum, 1× MEM non-essential amino acids solution, 1 mM sodium pyruvate, and 1% penicillin-streptomycin. AHA medium was replenished every second day. For the inhibition of secretion, Exo-1 (Sigma-Aldrich) was used at a working concentration of 100 nM in DMSO, which was replenished daily. Blockade of collagen VI-integrin interactions was performed using monoclonal antibodies against integrin β1 (10 μg/mL anti-integrin β1, Millipore sigma, MAB2253), Cilengitide (5 μg/mL) or soluble RGD peptides (sol RGD, 0.5 mM), added daily to culture media. To provide hypoxic conditions, cell-laden hydrogels were maintained in a hypoxic chamber (Biospherix, C-Chamber Incubator Subchamber) with a mixture of 1% O_2_, 5% CO_2_, and 94% N2 placed in a 37 °C humidified incubator. For normoxic conditions, cell-laden hydrogels were cultivated at 21% O_2_ in a 37 °C humidified incubator buffered with 5% CO_2_ as a comparison.

### Metabolic labeling for visualizing protein secretion

To visualize cell membranes and secreted proteins, hydrogels at the desired time point were rinsed with 1% bovine serum albumin (BSA, Sigma-Aldrich) in PBS and incubated in 10 μM AZDye 488 DBCO (Click Chemistry Tools) for 30 min at 37 °C, 5% CO2. After washing with 1% BSA-PBS, hydrogels were fixed in 4% paraformaldehyde solution for 30 min at room temperature followed by three washes in PBS. 1 μg/mL of wheat germ agglutinin (WGA, Invitrogen, Thermo Fisher Scientific) solution and 1 μg/mL of DAPI (Sigma-Aldrich) solution were incubated for 30 min at room temperature to stain the cell membrane and nucleus, respectively.

For staining of specific ECM molecules with metabolic labeling, the permeabilization step was added after hydrogel fixation by incubating in 0.2 % Triton X-100 in PBS for 2 hr at room temperature. Hydrogels were blocked with 2% BSA-PBS solution for 1 hr at room temperature and incubated with one or more of the following primary antibodies diluted in 2% BSA-PBS at 4 °C for 48∼72 hr: rabbit anti-Collagen VI (1:200; Abcam, ab182744), rabbit anti-fibronectin (1:200; Abcam, ab2413), and mouse anti-Tenascin C (1:200; Abcam, ab3970). After washing with 1% BSA-PBS, the samples were stained with goat anti-rabbit immunoglobulin (IgG) [heavy and light chains (H + L)] secondary antibody, Alexa Fluor 555 (Invitrogen) or goat anti-mouse immunoglobulin (IgG) [heavy and light chains (H + L)] secondary antibody, Alexa Fluor 594 (Invitrogen) for 2 hr at room temperature. Hydrogels were washed three times followed by DAPI staining (1:1,000; Invitrogen, D1306, in PBS) for 30 min at room temperature and stored at 4 °C until analyzed.

### Quantification of protein secretion

3D cell-laden hydrogel samples were imaged via confocal microscopy (LSM 880, Zeiss). All taken images were analyzed and post-processed with ImageJ for quantifying protein secretion by the cells. Briefly, each image channel z-stack was converted into a 16-bit grayscale image. After generating thresholded images to define the fluorescent signals, the integrated intensities from ECM and nucleus were measured. Normalized ECM secretion levels were determined by calculating the ratio of integrated intensities of ECM and nuclei.

### AFM measurement for hydrogel stiffness

Measurements of elastic moduli of cell-laden hydrogels were obtained as previously described (*61*) using an MFP-3D-Bio AFM (Asylum Research, Oxford Instruments). Cantilevers with colloidal probes were used to ensure large and well-defined contact areas with minimal tip-sample pressure. Briefly, polystyrene microspheres (diameter 25 μm; Polysciences, Inc) were mounted onto tipless triangular silicon nitride cantilevers (PNP-TR-TL-Au, spring constant (k) ≈ 0.08 N/m; Nanoworld) using UV-curable adhesive (Loctite, HL88489). Prior to sample measurements, cantilevers were calibrated by the thermal method. Cell-laden gels were taken directly from the cell culture incubator, rinsed, and immersed with clean PBS. Acellular or cell-laden hydrogels were transferred and adhered to a 50 mm Petri dish (Falcon, 351006) coated with 1 mg/mL poly-D-lysine (Sigma-Aldrich). Hydrogels were kept hydrated in PBS at room temperature throughout all processes and measured within 1 hr. Force measurements were obtained at randomly selected locations and performed at cell-hydrogel interfaces. Indentations were carried out with a relative trigger force of 1 nN and a loading rate of 1 μm/s, and force curves were taken in multiple 10 x 10 grids 10 um in size for each sample. E was extracted from force curves using the system software by fitting the Hertz model for a spherical tip of radius 12.5 μm.

### 3D Tumorsphere assays

GBM cells were harvested, dissociated, and plated at a density of 2.4×10^5^ cells/well in AggreWell™400 (StemCell™ Technologies) to create tumorspheres. After incubating the plate at 37 °C, 5% CO_2_ for 3 days, the desired number of tumorspheres were gently harvested in phenol-free DMEM (Gibco) and added to the HA mixture before cross-linking. The cell growth medium was lastly added and hydrogels were inspected by phase contrast microscopy (Eclipse TE2000-E, Nikon). Tumorspheres were tracked for up to 7 days, and invasion was quantified using ImageJ by normalizing the tumorsphere area to its area on ‘day 0’.

### Preparation of proteomic analysis samples

A previously described digestion scheme was used to collect secreted ECM proteins from the hydrogels in situ (*62*). Cell-laden gels were treated with decellularization solution (0.5% Triton X-100 supplemented with 0.5% NH_4_OH in distilled water) for 1 hr on a rotating shaker (100 rpm). Decellularized gels were then washed with PBS on a rotating shaker three times and stored in PBS at 4°C until analysis. Hydrogels were incubated for 10 min at room temperature with 100 μL acetonitrile (Optima, LC/MS Grade, Fisher Chemical) and then transferred to a 2 mL microcentrifuge tube. After removing the acetonitrile from the gels, 50 ng trypsin (Sequencing Grade Modified Trypsin, Lyophilized, Promega) was added to the hydrogels and incubated for 30 min to permeate the hydrogels at room temperature. 100 μl of 100 mM ammonium bicarbonate was then added, and the hydrogel was incubated overnight at 37 °C on a benchtop mixer with shaking at 800 rpm. The tube was centrifuged (425 xg rcf, 3 min, 25 °C), and the supernatant was transferred to a new 2 mL microcentrifuge tube. The hydrogel was then incubated with 30 μl acetonitrile for 10 min at room temperature. Finally, the tube was centrifuged again (425 xg rcf, 3 min, 25 °C), and this supernatant was combined with the previous supernatant to yield the final peptide sample.

### Mass spectrometry and proteomic data analysis

Trypsin-digested samples were submitted to the Vincent J. Coates Proteomics/Mass Spectrometry Laboratory, UC Berkeley for analysis. Peptides were loaded to a nanoscale HPLC column composed of 10 cm of Polaris C18 5 μm packing material (Agilent), followed by 4 cm of Partisphere 5 SCX (Whatman). The column was then directly coupled to an electrospray ionization source mounted on a LTQ XL linear ion trap mass spectrometer (Thermo Fisher). Peptides were eluted using an 8-step multidimensional protein identification technology (MudPIT) procedure (*63*). Protein identification and quantification were performed with Integrated Proteomics Pipeline (IP2, Integrated Proteomics Applications, Inc. San Diego, CA). Tandem mass spectra extracted using RawExtractor (*68*) were searched against the human protein databases downloaded from NCBI supplemented with sequences of common contaminants, concatenated to a decoy database in which the sequence for each entry in the original database was reversed (*69*). Spectra data was searched with 3000.0 milli-amu precursor tolerance and the fragment ions were restricted to a 600 ppm tolerance. Search space included all fully tryptic peptide candidates with no missed cleavage restrictions. Cysteine carbamidomethylation (+57.02146) was considered a static modification. We required 1 peptide per protein and both tryptic termini for each peptide identification. The ProLuCID search results were assembled and filtered using the DTASelect program (*64–66*) with a peptide false discovery rate (FDR) of 0.001 for single peptides and a peptide FDR of 0.005 for additional peptides for the same protein. The normalized spectral abundance factor (NSAF) was used for label-free quantification, which is defined as a ratio of the spectral counts for a given protein to its length. Identified proteins from the hydrogels were compared with the datasets of Matrisome Project (matrisomeproject.mit.edu/proteins) to annotate corresponding matrisomes.

### Transcriptomic analysis

We obtained human glioma transcriptomic data from the Rembrandt and The Cancer Genome Atlas (TCGA) patient databases via the GlioVis data portal (http://gliovis.bioinfo.cnio.es/). Using both databases, we identified highly expressed genes in GBM tissue, compared to non-tumor using adjusted p-value under 0.01 and log_2_(fold change) over 0.25. Statistical significance was assessed by performing two-tailed unpaired Student’s t-test with equal variances, and the resulting p-values were adjusted based on Benjamini-Hochberg correction. Genes identified from Rembrandt and TCGA in common were then compared against human MatrisomeDB (*36*) to be classified into core ECM or ECM-associated. Gene Ontology (GO) enrichment analysis (http://geneontology.org) was performed for upregulated genes in GBM compared to non-tumor in terms of the molecular function (MF), biological process (BP), and cellular component (CC). The above-mentioned analysis was repeated by comparing GBM tissue and low-grade glioma using both databases. Commonly upregulated genes in GBM tissues compared to non-tumor and low-grade glioma were defined as GBM-enriched genes. The Ivy GAP database (http://glioblastoma.alleninstitute.org/) was used to analyze the region-specific transcriptome profiles of GBM-enriched genes. We made an expression heatmap of GBM-enriched genes using Morpheus software (https://software.broadinstitute.org/morpheus).

### Real-time quantitative polymerase chain reaction (qPCR)

Hyaluronidase (750-3000 units/mg, Sigma-Aldrich, H3884) was added to cell-laden HA hydrogels and pipetted to physically dissociate gels. After incubating at 37 °C for 1 hr, the degraded HA containing the cells was centrifuged to collect the cell pellet. Total mRNA was then isolated by using RNeasy Micro kit (Qiagen) according to the manufacturer’s protocol, followed by cDNA synthesis using Iscript^TM^ cDNA Synthesis Kit (Bio-Rad, 1708891). qRT-PCR was performed using 2x SYBR qPCR Master Mix (Bimake, B21203) for 40 cycles in a CFX real-time PCR detection system (Bio-Rad). To quantify relative fold change in the level of genes, the qRT-PCR data were analyzed by calculating ΔΔCt with respect to GAPDH. The primer sequences used in this study are listed in Supplementary Table 1.

### Proximity Ligation Assay

Duolink® proximity ligation assays (PLA, Sigma-Aldrich) were performed to visualize colocalization between collagen VI and integrins, following the manufacturer’s protocol. Briefly, the cell-laden hydrogels were fixed using 4% paraformaldehyde (PFA) for 30 min and permeabilized with 0.2 % Triton-X solution for 2 hr. Primary antibodies for collagen VI (Abcam, ab182744) and integrins (Sigma-Aldrich, MAB2253 for anti-integrin β1, Abcam, ab190147 for anti-integrin αvβ3) were added at 1:100 dilution in 1% BSA-PBS solution at 4 °C for 48 ∼ 72 hr. After two 5-min washes in PBS, PLUS and MINUS PLA probes conjugated with oligonucleotides to tag the primary antibodies were incubated for 2 hr at 37 °C. Ligation solution was added to the sample and incubated for 1 hr at 37 °C after washing steps. After removing the ligation solution, the hydrogels were incubated with an amplification solution for 2 hr at 37 °C. Following final washes, the nucleus staining was performed with DAPI solution and stored at 4°C until imaging. Samples were imaged using confocal microscopy (LSM 880, Zeiss) with a collection of z-stacks and PLA quantification via ImageJ. Briefly, fluorescent signals from collagen VI-integrin complex were projected into z-stack and converted into 16-bit grayscale. Thresholded images were converted to binary images, and the area of fluorescent regions was measured. The fluorescent PLA area was divided by the number of nuclei to calculate the collagen VI/integrin complexes per cell.

### Luciferase assay

A lentiviral construct encoding a 7xTFP T cell factor/lymphoid enhancer factor (TCF/LEF) luciferase reporter (*70*) was used to transduce U87 cells to measure activation of the β-catenin signaling pathway. Briefly, lentiviral particles were packaged with psPAX2 and pMD2.G in human embryonic kidney (HEK) 293 T cells via polyethylenimine (PEI) transfection. U87 cells were infected with purified viral particles with a multiplicity of infection (MOI) of 1. Infected cells were encapsulated within HA gels under each condition, and the cell pellets were collected by hyaluronidase treatment at the desired time point. After rinsing once with PBS, lysis reagent (Promega) was added to the cell pellet and then the pellet debris was removed by centrifugation. 20 μl of supernatants were transferred to each well of a white opaque 96-well plate, and 100 μl of Luciferase Assay Reagent (Promega) was treated right before detection. Luminescence intensity was measured via SpectraMax luminometer (Molecular Devices). To compensate for different proliferation rates across conditions, total protein concentration was additionally measured through BCA protein assay (Pierce) and used to normalize the luminescence signals.

### Immunostaining and quantification of β-catenin nuclear translocation

Hydrogels at the desired time point were rinsed with 1% bovine serum albumin (BSA, Sigma-Aldrich) in PBS and fixed in 4% paraformaldehyde solution for 30 min at room temperature followed by three washes in PBS. Fixed hydrogels were permeabilized with 0.2 % Triton X-100 and incubated for 2 hr at room temperature. Hydrogels were then blocked in 2% BSA-PBS solution for 1 hr at room temperature and then incubated with a primary antibody against β-catenin (1:100; Cell signaling technology, #9562) diluted in 2% BSA-PBS, and stored at 4 °C for 48∼72 hr. After washing with 1% BSA-PBS, the samples were stained with Goat anti-Rabbit IgG (H+L) Cross-Adsorbed Secondary Antibody, Alexa Fluor 488 (Invitrogen) for 2 hr at room temperature. Hydrogels were washed three times followed by staining for F-actin (1:1,000; Alexa Fluor™ 546 Phalloidin, Invitrogen), and nuclei (DAPI, 1:1,000; Invitrogen, D1306) in PBS for 30 min at room temperature and stored at 4 °C until imaged. Samples were visualized with a confocal microscope (LSM 880, Zeiss) with collection of z-stacks. β-catenin nuclear- to-cytosolic ratio was calculated by measuring fluorescence intensity using ImageJ. Briefly, binary masks of nuclei and F-actin were created to differentiate nuclear and cytosolic areas. Then, cytosol-only masks were created by overlaying nuclei and F-actin masks to exclude the nucleus. The integrated intensity of β-catenin fluorescence was quantified in the regions of the nucleus and cytosol-only masks, and divided by the area of each mask, resulting in nuclear and cytosolic β-catenin, respectively. β-catenin nuclear localization was determined by calculating the ratio between nuclear β-catenin and cytosolic β-catenin.

### Statistical analysis

Graphical representation and statistical analyses of the data from this study were generated using GraphPad Prism 9. The data were presented as the means ± S.D. The unpaired, two-sided Student’s t-tests and one-way or two-way analysis of variance (ANOVA) followed by Tukey’s multiple comparisons test were conducted to determine the statistical significance of data between groups. Throughout the study, one experiment was performed with at least three samples, and one to three independent experiments were conducted to ensure the reproducibility of the results. The number of biological replicates was the sum of sample replicates collected from independent experiments, as described in the appropriate figure legends.

## Acknowledgments

Proteomic studies were performed at the Vincent J. Coates Proteomics/Mass Spectrometry Laboratory at UC Berkeley, supported in part by NIH S10 Instrumentation Grant S10RR025622. We thank Dr. Lori Kohlstaedt (Vincent J. Coates Proteomics/Mass Spectrometry Laboratory, UC Berkeley) for contributing methodological descriptions of mass spectrometry and proteomic data analysis. Glioma stem cell lines were developed at the University of Texas M. D. Anderson Department of Neurosurgery with support by grants from the National Cancer Institute (1R01CA214749, 1R01CA247970, P30CA016672 and 2P50CA127001) and the University of Texas M. D. Anderson Moon Shots Program^TM^. The 7xTFP T cell factor/lymphoid enhancer factor (TCF/LEF) luciferase reporter construct was obtained from Dr. David V. Schaffer’s laboratory at UC Berkeley.

## Author contributions

J.C. designed and performed the experiments, analyzed, and interpreted the data, and wrote the manuscript. E.A.D. contributed to conducting and completing the experiment regarding AFM measurement on 3D cell-laden hydrogels. E.M.C. contributed to the synthesis and characterization of HA hydrogels. A.F. performed AFM experiments and data validation.

M.K.A. and S.K. supervised the project, helped in the experimental design and data interpretation, and wrote the manuscript.

## Data availability

All data needed to evaluate the conclusions in the paper are present in the paper and/or the Supplementary Materials. Additional data related to this paper may be requested from the authors.

