## Supplementary Information for "Glioma Cells Secrete Collagen VI to Facilitate Invasion"

**Supplementary Methods**

**3D cell encapsulation within Collagen I and Matrigel**

Collagen I (high concentration, rat tail, Corning, 354249) and Matrigel (Corning, 354230) solutions were held on ice until immediately before use and assembled into gels via temperature switch following standard protocols. Briefly, a high-concentration stock of Collagen I was diluted to 4.0 mg/mL in distilled water. GBM cells were dissociated and counted at the desired concentration in PBS and added to this diluted collagen I solution, and then 1 M NaOH was added to bring the pH to 7.4 for cell culture. The Matrigel stock solution was diluted to 4.0 mg/mL with cell culture medium containing the cells at the desired concentration. Solutions were mixed thoroughly before gelation. For both materials, hydrogel assembly was induced by incubation at 37 °C for 1 hr while immersed in cell culture medium. After a 3-day incubation, hydrogels were rinsed with 1% bovine serum albumin (BSA, Sigma-Aldrich) in PBS and fixed in 4% paraformaldehyde solution for 30 min at room temperature. Hydrogels were washed three times, fixed, and stained for F-actin (1:1,000; Alexa Fluor™ 546 Phalloidin, Invitrogen) and nuclear DNA (DAPI, 1:1,000; Invitrogen, D1306) in PBS for 30 min at room temperature and stored at 4 °C until imaged. Samples were imaged using a confocal microscope (LSM 880, Zeiss) and visualized by z-stack mode. Image post-processing was conducted using ImageJ.

**Characterization of decellularized hydrogels**

Gels were chemically decellularized (0.5% Triton X-100 supplemented with 0.5% $NH_4OH$ in distilled water) and examined using the Eclipse TE2000-E microscope (Nikon). HA hydrogels before and after decellularization were treated with hyaluronidase (750-3000 units/mg, Sigma-

Aldrich, H3884) to collect the cells before DNA extraction. DNA content before and after decellularization was determined by extracting DNA from each hydrogel using QIAamp® DNA Micro Kit (Qiagen) and measuring DNA concentration with a spectrophotometer. For quantification of peptide concentration after decellularization, Pierce™ quantitative colorimetric peptide assay kit (Thermo Fisher Scientific, 21013024) was used according to the manufacturer's protocol.

**DIC imaging**

Cell-containing HA mixtures were plated on glass bottom 6-well plates (MatTek Corporation, P06G-1.5-20-F), and cell culture medium was added after crosslinking. DIC images were taken using an Eclipse TE2000 Microscope (Nikon) equipped with a polarizer, prisms, and analyzer using a Nikon 60x/1.40 Oil Plan Apo VC DIC N2 objective. Any cells located at the hydrogel edge or contacting the glass surface were excluded from the imaging.

**Collagen VI functional studies**

U87 $Bev^S$ or $Bev^R$ cells were engineered for genetic alteration of COL6A3 using CRISPR Gene Knockout Kit v2 (Synthego). Briefly, target-specific multi-guide sgRNA sequences were designed for COL6A3, including sequences GAUCGCUUUCGACUCCUCCC, GCAUAAUGGCCUUUGCCAUU, GUGAUCAGCACCAAAAGCUU. TruGuide™ sgRNA non-targeting negative control (Invitrogen) was used as a control. Multi-guide sgRNA and Cas9 2NLS nuclease (Synthego) were mixed at a ratio of 1.3:1 in Opti-MEM™ I Reduced Serum Medium (Thermo Fisher Scientific) to create ribonucleoprotein (RNP). RNPs were combined with Lipofectamine™ CRISPRMAX™ Reagent in Opti-MEM™ and then mixed with the cells before seeding into the plates. After 72 h of lipofection, limiting dilution was performed to clonally expand the edited cell population. KO clones were expanded and screened for expression of the target gene and protein by qPCR and collagen VI immunostaining as confirmation of the KO cells used for further study.

To further study the effect of exogenous collagen VI proteins, native human Collagen VI protein (Abcam, ab7538) was used. Collagen VI was added to the growth media proteins at a concentration of 20 μg/ml, and Collagen VI-supplemented growth media was replenished every 2 days.

**Blockade of β-catenin signaling**

To examine the effect of β-catenin activity on GBM invasion, two inhibitors to perturb β-catenin signaling were added to the cell culture medium during the 3D tumorsphere assay. 50 μM of iCRT14 (Selleck Chemicals) was used to inhibit the β-catenin-TCF/LEF interaction, and 300 nM of Laduviglusib (CHIR-99021, Selleck Chemicals) was used as a highly potent and specific glycogen synthase kinase-3 (GSK3) inhibitor that potentiates β-catenin activity during 3D invasion. HA hydrogels were immersed with an inhibitor-containing cell culture medium and media was replenished daily.

**GSC culture**

All GSCs were propagated as neurospheres at 37 °C, 5% $CO_2$ in DMEM/F12 basal medium supplemented with 2 v/v% B-27 supplement (Gibco), 20 ng/mL EGF (R&D Systems), and 20 ng/mL FGF (R&D Systems). GSCs were dissociated into single cells using Accutase (Innovative Cell Technologies). Cells used in this study were passaged <30 times, screened for mycoplasma, and authenticated every six months by Short Tandem Repeat (STR) analysis at the University of California Cell Culture Facility. For visualization of secreted ECM, cell-laden hydrogels were cultured in GSC AHA growth media, which consists of 1 : 1 ratio mixture of DMEM, high glucose, no glutamine, no methionine, no cystine (Gibco) and Ham's F-12 Nutrient (Gibco) as a basal medium with 2 v/v% B-27 supplement (Gibco), 20 ng/mL EGF (R&D Systems), and 20 ng/mL FGF (R&D Systems), supplemented with 0.2 mM AHA (Click chemistry tools), 2.5 mM L-glutamine, and 0.1 mM cystine (Sigma-aldrich).

**Supplementary Tables**

**Table 1. qPCR primers**

| **Target** | **Forward (5'-3')** | **Reverse (5'-3')** |
|---|---|---|
| GAPDH | GCATCCTGGGCTACACTGAG | GTCAAAGGTGGAGGAGTGGG |
| COLVI | GTTTTGTTCTTAAAGGCCCTGC | CAGACACTGTGAGGCCTGAA |
| ZEB1 | AAGAATTCACAGTGGAGAGAAGCCA | CGTTTCTTGCAGTTTGGGCATT |
| CD44 | CGAAGAAGGTGTGGGCAGAA | CCGTTGAGTCCACTTGGCTT |
| MMP2 | AGCGAGTGGATGCCGCCTTTAA | CATTCCAGGCATCTGCGATGAG |
| MMP9 | TTGACAGCGACAAGAAGTGG | GCCATTCACGTCGTCCTTAT |
| ITGB1 | CAAGAGAGCTGAAGACTATCCCA | TGAAGTCCGAAGTAATCCTCCT |

### Supplementary Figures

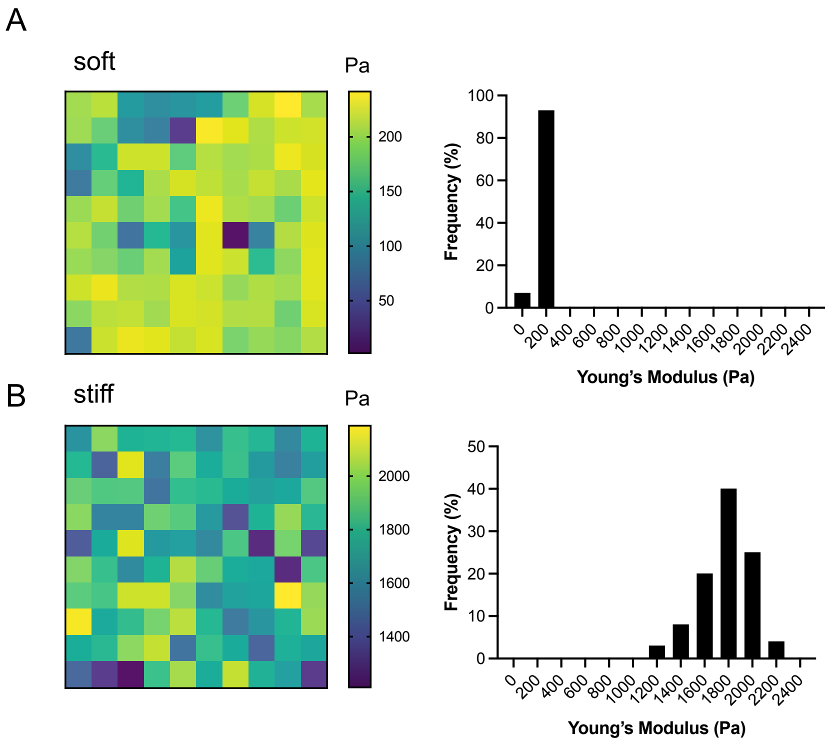


**Fig. S1. Measurement of the HA hydrogel stiffness using AFM**. (**A**) Representative force map (left) and stiffness distribution (right) of acellular 'soft' HA/RGD-. (**B**) Representative force map (left) and stiffness distribution (right) of acellular 'stiff' HA/RGD-.

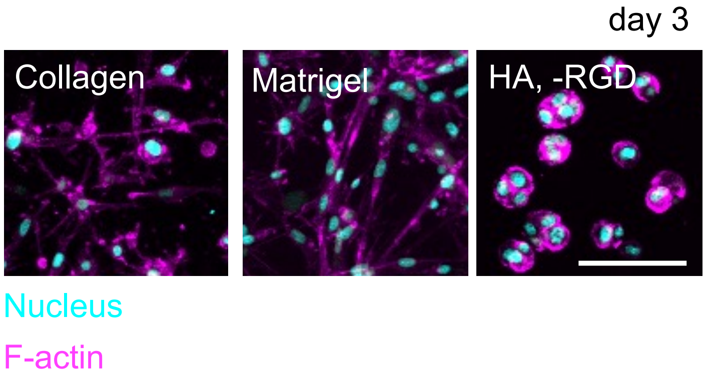


**Fig. S2. 3D encapsulation of U87 GBM cells in collagen type I, Matrigel, and HA/RGD- gels.** Representative fluorescence images of U87 cells after 3-day cultivation. Cyan: nucleus, Magenta: F-actin. Scale bar: 100 µm.

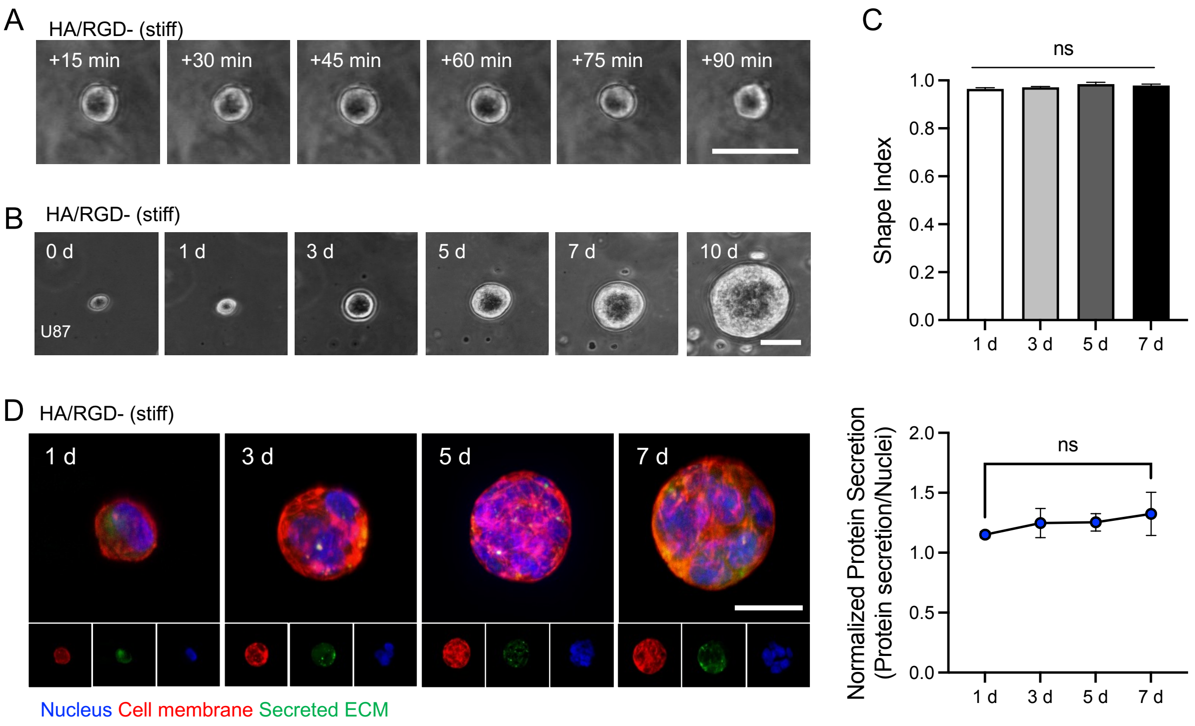


**Fig. S3**. **Matrix remodeling of GBM cells within 3D stiff HA hydrogels.** (**A**) Short-term time-lapse observation of U87 GBM cells interacting with stiff HA/RGD- gels. Scale bar: 50 μm. (**B**) Representative time course of 3D encapsulated U87 GBM cells within stiff HA/RGD- gels for 10 days. Scale bar: 50 μm. (**C**) Evolution of GBM cell shape index after stiff HA/RGD- encapsulation. (**D**) Representative images of proteins secreted by GBM cells within stiff HA/RGD- gels over time via metabolic labeling (left). Quantification of fluorescence signal from secreted proteins by GBM cells, normalized by the intensities of nuclei (right). Statistical significance was analyzed using one-way ANOVA followed by Tukey's multiple comparisons test. ns: no significance.

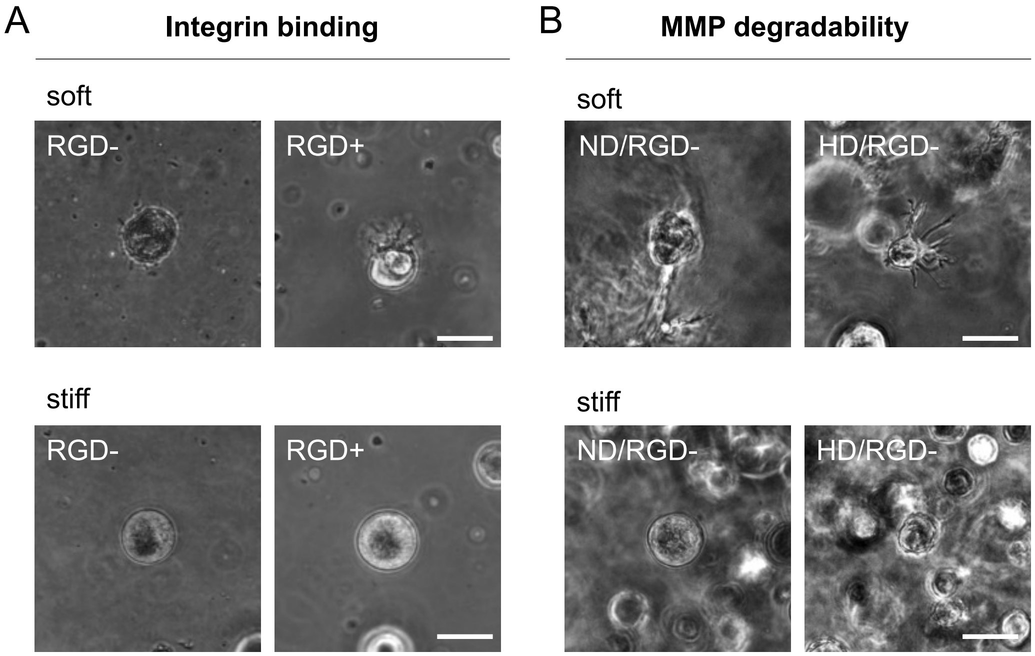


**Fig. S4**. **Dependence of 3D U87 GBM cell morphology on HA gel properties.** Representative phase contrast images of GBM cells within HA gels as a function of (**A**) integrin ligation (RGD- vs RGD+) and (**B**) protease degradability (ND/RGD- vs HD/RGD-). Scale bar: 50 µm.

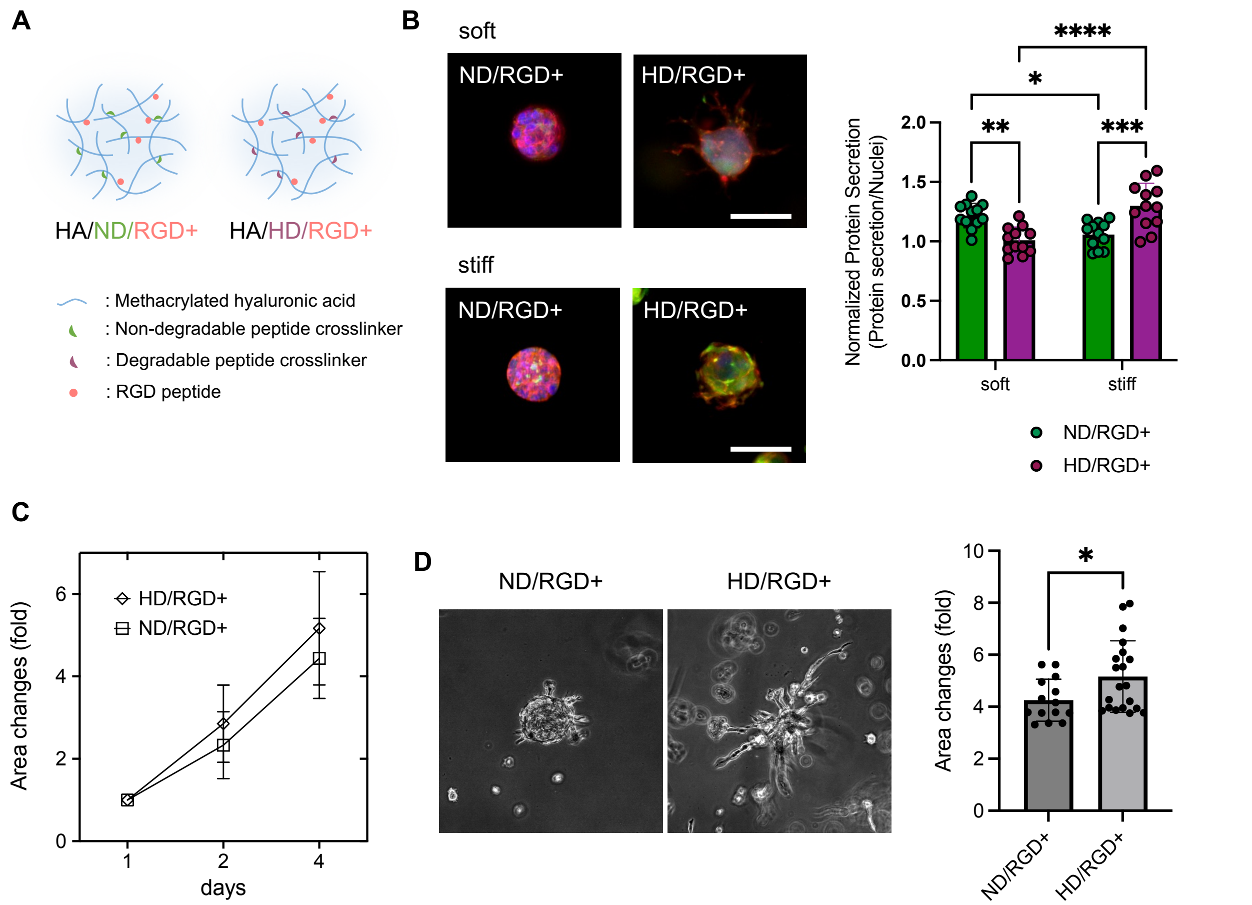


**Fig. S5. ECM remodeling by GBM cells within HA hydrogels featuring both RGD peptides and MMP-cleavable crosslinkers.** (**A**) Schematic illustration of a set of HA/RGD+ models depending on degradability (ND and HD). (**B**) Representative images of fluorescently labeled proteins secreted by GBM cells in integrin-ligating matrices of differing MMP degradability (ND/RGD+ vs HD/RGD+). Scale bar: 50 µm. (left) Quantification of fluorescent signals from secreted proteins by GBM cells, normalized by the number of nuclei. (right) (n = 12) (**C**) Quantification of 3D GBM tumorsphere invasion in these matrices over 4 days. (n = 14 to 20) (**D**) Representative images (right) and quantification (left) of GBM tumorsphere invasion at day 4. Statistical significance was analyzed using a two-way ANOVA with Tukey's multiple comparisons test (**B**) or an unpaired two-sided Student's t-test (**D**). ****$p < 0.0001$, ***$p < 0.001$, **$p < 0.01$, *$p < 0.05$.

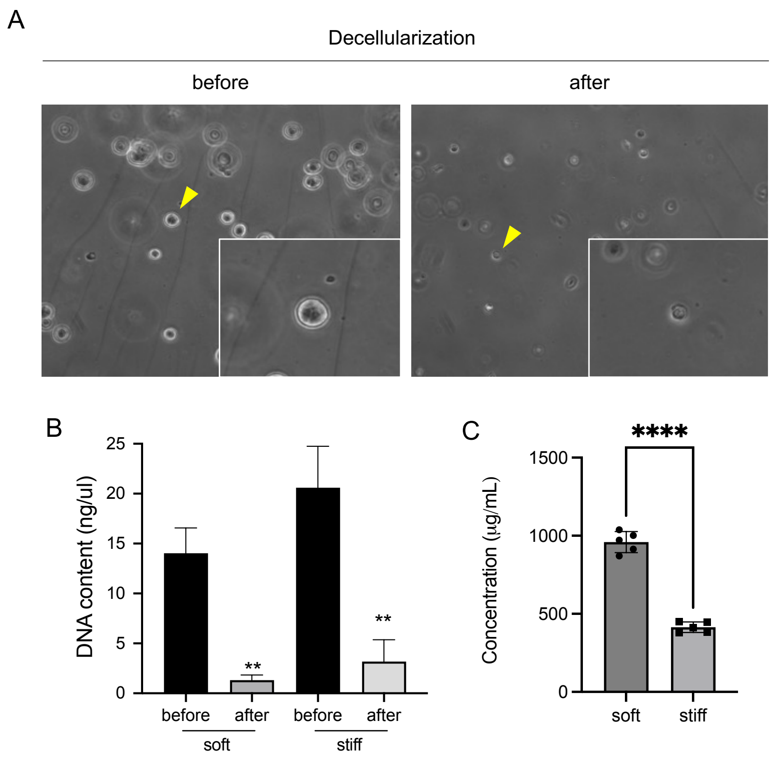


**Fig. S6**. **Characterization of decellularized hydrogels.** (**A**) Representative images of HA/RGD- gels before and after decellularization. The yellow arrows indicate the insets within the figure. (**B**) Measurement of DNA content for 3D cell-laden soft and stiff HA/RGD- hydrogels before and after decellularization. (n = 3) (**C**) Peptide concentration of decellularized soft and stiff HA/RGD- gels. (n=3) Statistical significance was analyzed using an unpaired two-sided Student's t-test (**B**, **C**). ****$p < 0.001$, **$p < 0.01$.

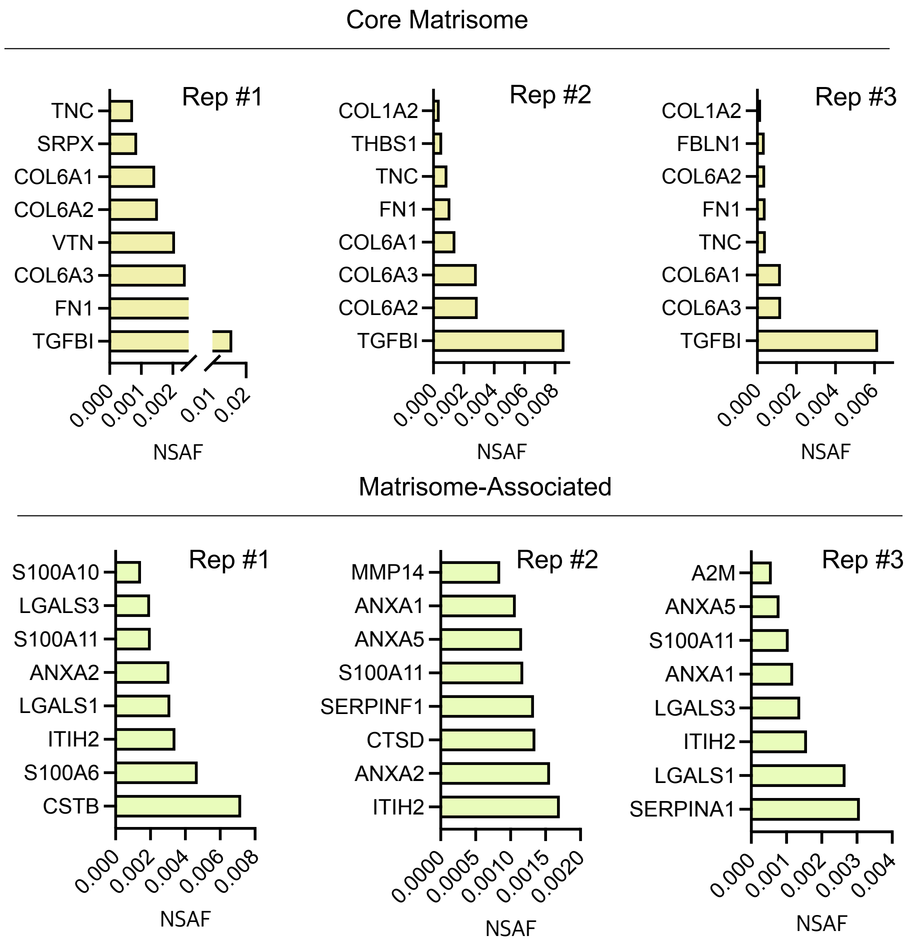


**Fig. S7.** Comparison of the top eight core matrisome and matrisome-associated proteins with the highest expression in HA/RGD- samples derived from different experimental replicates (Rep #1-3). NSAF: normalized spectral abundance factor.

**Detected Matrisome Proteins depending on HA gel properties**

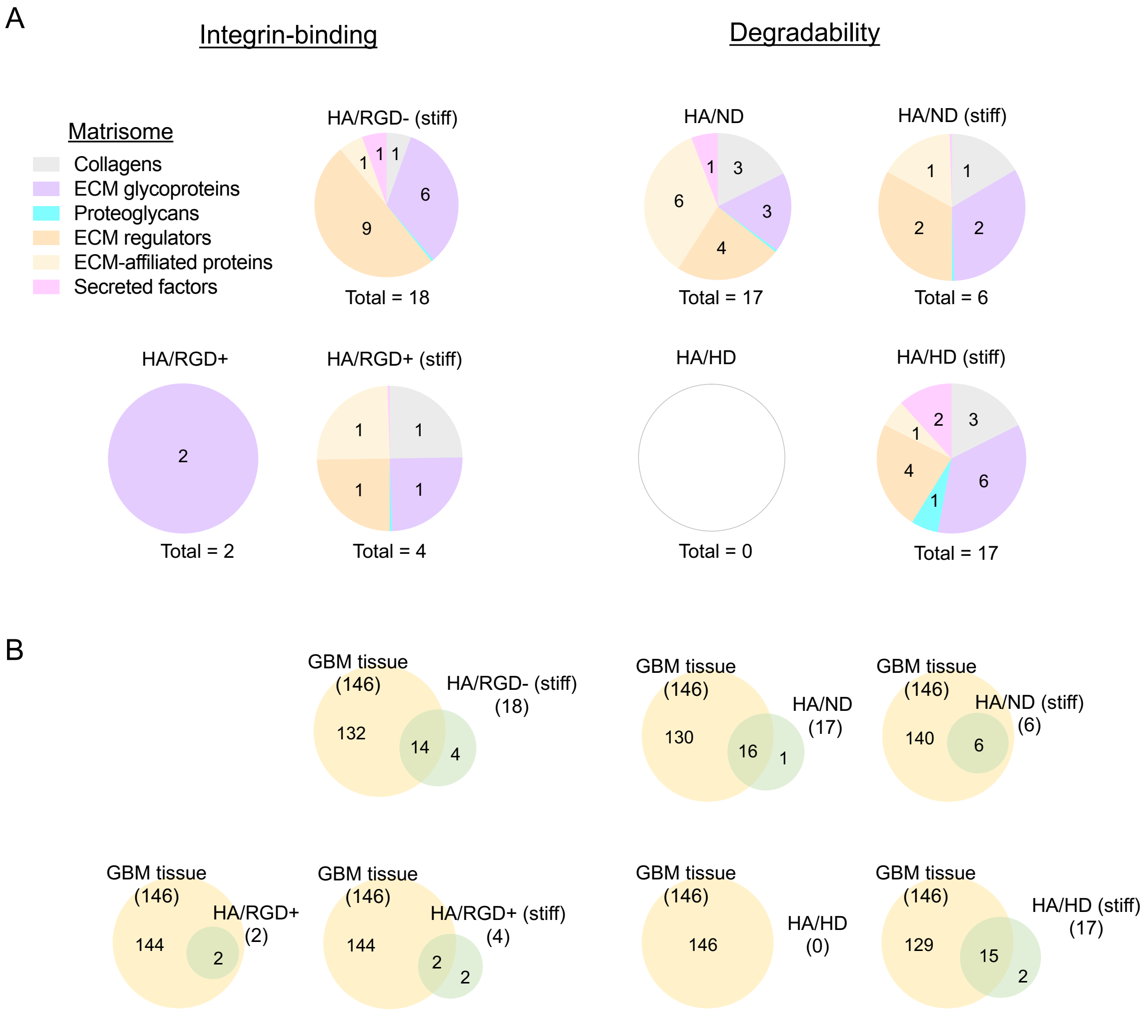


| | | Integrin binding | | | | MMP degradability | | | |
|---|---|---|---|---|---|---|---|---|---|
| Category | GBM tissue (%) | HA/RGD- | HA/RGD- (stiff) | HA/RGD+ | HA/RGD+ (stiff) | HA/ND | HA/ND (stiff) | HA/HD | HA/HD (stiff) |
| Collagens | 12.33 | 20.00 | 5.56 | 0.00 | 25.00 | 17.65 | 16.67 | - | 17.65 |
| ECM glycoproteins | 30.82 | 20.00 | 33.33 | 100.00 | 25.00 | 17.65 | 33.33 | - | 35.29 |
| Proteoglycans | 7.53 | 2.86 | 0.00 | 0.00 | 0.00 | 0.00 | 0.00 | - | 5.88 |
| ECM regulators | 26.71 | 34.29 | 50.00 | 0.00 | 25.00 | 23.53 | 33.33 | - | 23.53 |
| ECM-affiliated proteins | 13.70 | 14.29 | 5.56 | 0.00 | 25.00 | 35.29 | 16.67 | - | 5.88 |
| Secreted factors | 8.22 | 8.57 | 5.56 | 0.00 | 0.00 | 5.88 | 0.00 | - | 11.76 |

**Fig. S8. ECM compositions secreted by GBM cells highly depend on HA gel properties.** (**A**) Pie charts showing the composition of matrisome proteins detected in decellularized HA gels of different integrin-ligation and degradability status. (**B**) Venn diagrams showing overlap of matrisome proteins secreted in various HA matrices and GBM tissue. The table shows more detailed category comparisons.

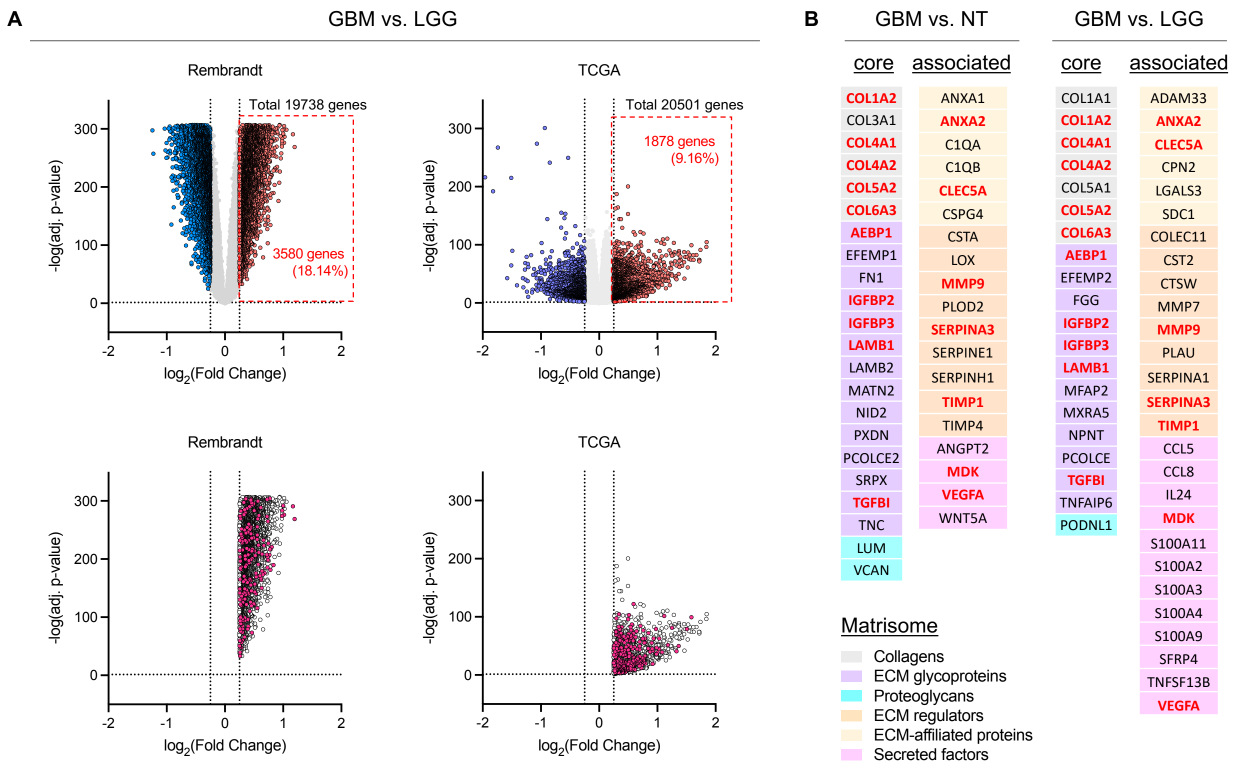


**Fig. S9. Identification of genes enriched in GBM tissues over low-grade glioma (LGG) tissues using Rembrandt and TCGA databases.** (**A**) (Upper row) Volcano plots from Rembrandt and TCGA to identify genes elevated in GBM compared to LGG. (lower row) Decomposition of upregulated genes into non-matrisomal (open circles) and matrisomal genes (purple circles). (**B**) Lists of enriched matrisomal genes in GBM vs. NT (left) and LGG (right). Commonly upregulated genes are indicated in red.

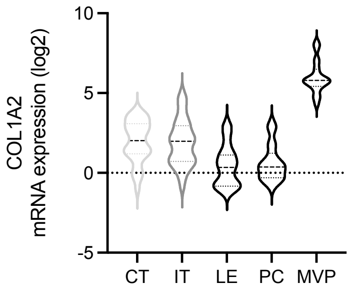


| Tukey's multiple comparisons test | Mean Diff. | 95.00% CI of diff. | Below threshold? | Summary | Adjusted P Value |
|---|---|---|---|---|---|
| CT vs. IT | 0.07086 | -0.8714 to 1.013 | No | ns | 0.9996 |
| CT vs. LE | 1.504 | 0.4948 to 2.512 | Yes | *** | 0.0006 |
| CT vs. PC | 1.333 | 0.3906 to 2.275 | Yes | ** | 0.0014 |
| CT vs. MP | -4.029 | -4.961 to -3.098 | Yes | **** | <0.0001 |
| IT vs. LE | 1.433 | 0.3761 to 2.489 | Yes | ** | 0.0025 |
| IT vs. PC | 1.262 | 0.2688 to 2.255 | Yes | ** | 0.0055 |
| IT vs. MP | -4.100 | -5.083 to -3.117 | Yes | **** | <0.0001 |
| LE vs. PC | -0.1707 | -1.227 to 0.8859 | No | ns | 0.9916 |
| LE vs. MP | -5.533 | -6.580 to -4.486 | Yes | **** | <0.0001 |
| PC vs. MP | -5.362 | -6.345 to -4.379 | Yes | **** | <0.0001 |

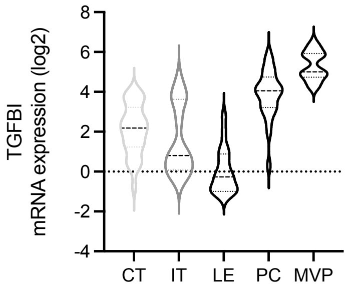


| Tukey's multiple comparisons test | Mean Diff. | 95.00% CI of diff. | Below threshold? | Summary | Adjusted P Value |
|---|---|---|---|---|---|
| CT vs. IT | 0.5151 | -0.4327 to 1.463 | No | ns | 0.5607 |
| CT vs. LE | 2.097 | 1.083 to 3.112 | Yes | **** | <0.0001 |
| CT vs. PC | -1.763 | -2.711 to -0.8153 | Yes | **** | <0.0001 |
| CT vs. MP | -3.191 | -4.128 to -2.254 | Yes | **** | <0.0001 |
| IT vs. LE | 1.582 | 0.5194 to 2.645 | Yes | *** | 0.0007 |
| IT vs. PC | -2.278 | -3.277 to -1.279 | Yes | **** | <0.0001 |
| IT vs. MP | -3.706 | -4.695 to -2.717 | Yes | **** | <0.0001 |
| LE vs. PC | -3.860 | -4.923 to -2.798 | Yes | **** | <0.0001 |
| LE vs. MP | -5.288 | -6.341 to -4.235 | Yes | **** | <0.0001 |
| PC vs. MP | -1.428 | -2.417 to -0.4386 | Yes | ** | 0.0010 |

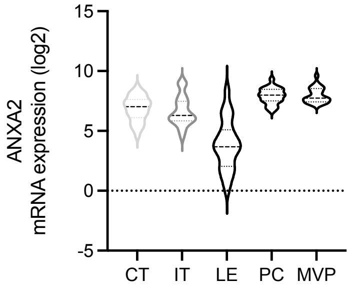


| Tukey's multiple comparisons test | Mean Diff. | 95.00% CI of diff. | Below threshold? | Summary | Adjusted P Value |
|---|---|---|---|---|---|
| CT vs. IT | 0.1918 | -0.7105 to 1.094 | No | ns | 0.9765 |
| CT vs. LE | 2.972 | 2.006 to 3.938 | Yes | **** | <0.0001 |
| CT vs. PC | -1.074 | -1.976 to -0.1717 | Yes | * | 0.0111 |
| CT vs. MP | -1.122 | -2.014 to -0.2300 | Yes | ** | 0.0061 |
| IT vs. LE | 2.780 | 1.769 to 3.792 | Yes | **** | <0.0001 |
| IT vs. PC | -1.266 | -2.217 to -0.3147 | Yes | ** | 0.0031 |
| IT vs. MP | -1.314 | -2.255 to -0.3725 | Yes | ** | 0.0017 |
| LE vs. PC | -4.046 | -5.058 to -3.035 | Yes | **** | <0.0001 |
| LE vs. MP | -4.094 | -5.097 to -3.092 | Yes | **** | <0.0001 |
| PC vs. MP | -0.04821 | -0.9897 to 0.8933 | No | ns | >0.9999 |

**Fig. S10**. **Expressions of COL1A2, TGFBI, and ANXA2 mRNA from Ivy_GAP in various histological regions**. (left) mRNA expression levels in various GBM subregions: Cellular tumor (CT), Invasive tumor (IT), Leading edge (LE), Pseudopalisading cells (PC), and Microvascular proliferation (MVP). (right) Summary tables of statistical comparisons between the histological regions. Statistical significance was analyzed using one-way ANOVA followed by Tukey's multiple comparisons test, ****$p < 0.0001$, ***$p < 0.001$, **$p < 0.01$, *$p < 0.05$, ns: no significance.

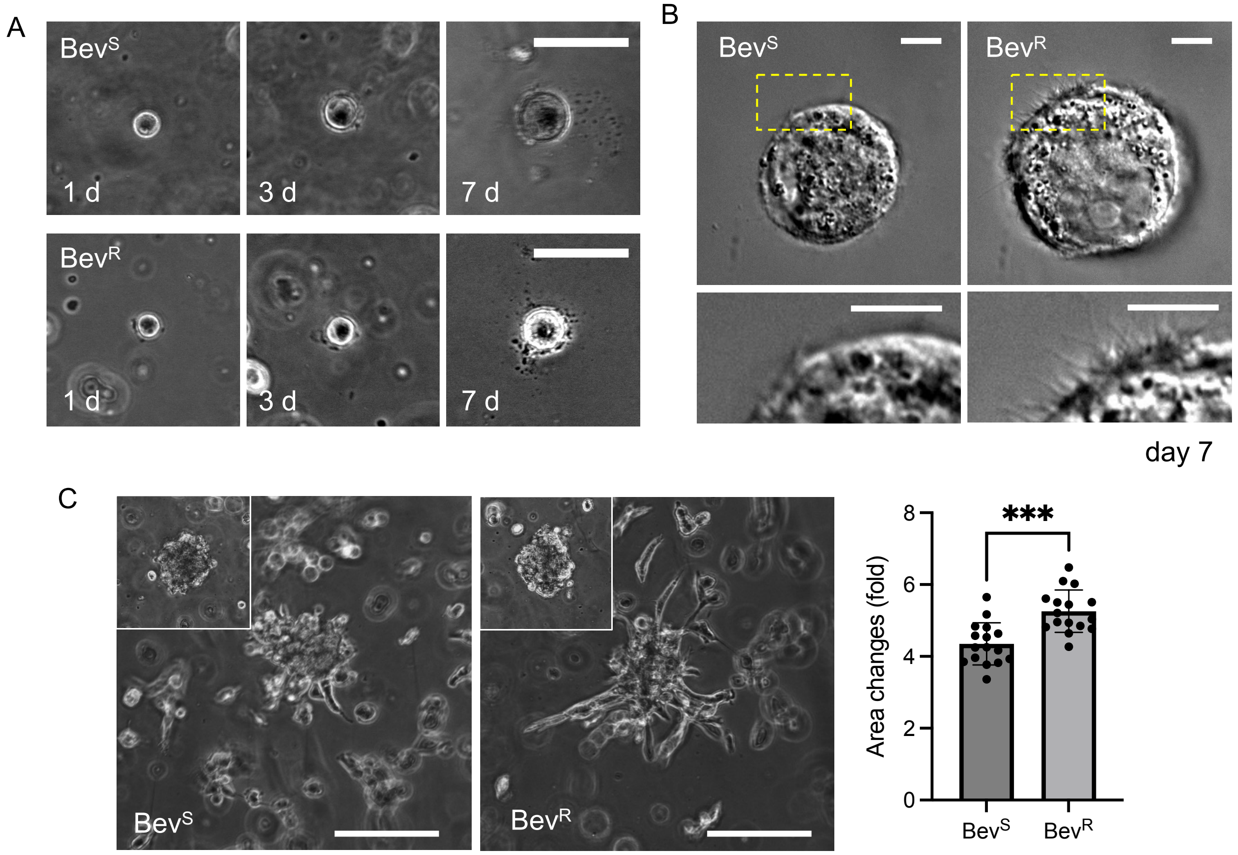


**Fig. S11. Comparison of Bev$^S$ and Bev$^R$ cell growth and invasion within 3D HA/RGD- gels.** (**A**) Representative phase images (left) of Bev$^S$ and Bev$^R$ cells within 3D HA/RGD- gels over 7 days of culture. Scale bar: 50 µm. (**B**) Representative differential interference contrast (DIC) images of Bev$^S$ and Bev$^R$ cells within 3D HA/RGD- gels at day 7. Scale bar: 50 µm. (**C**) Representative images (left) and quantification (right) of invasion of Bev$^S$ and Bev$^R$ tumorspheres within 3D HA/HD/RGD+ gels at day 4. (n = 16) Insets: tumorspheres at day 0. Scale bar: 100 µm. Statistical significance was analyzed using an unpaired two-sided Student's t-test. ***$p < 0.001$.

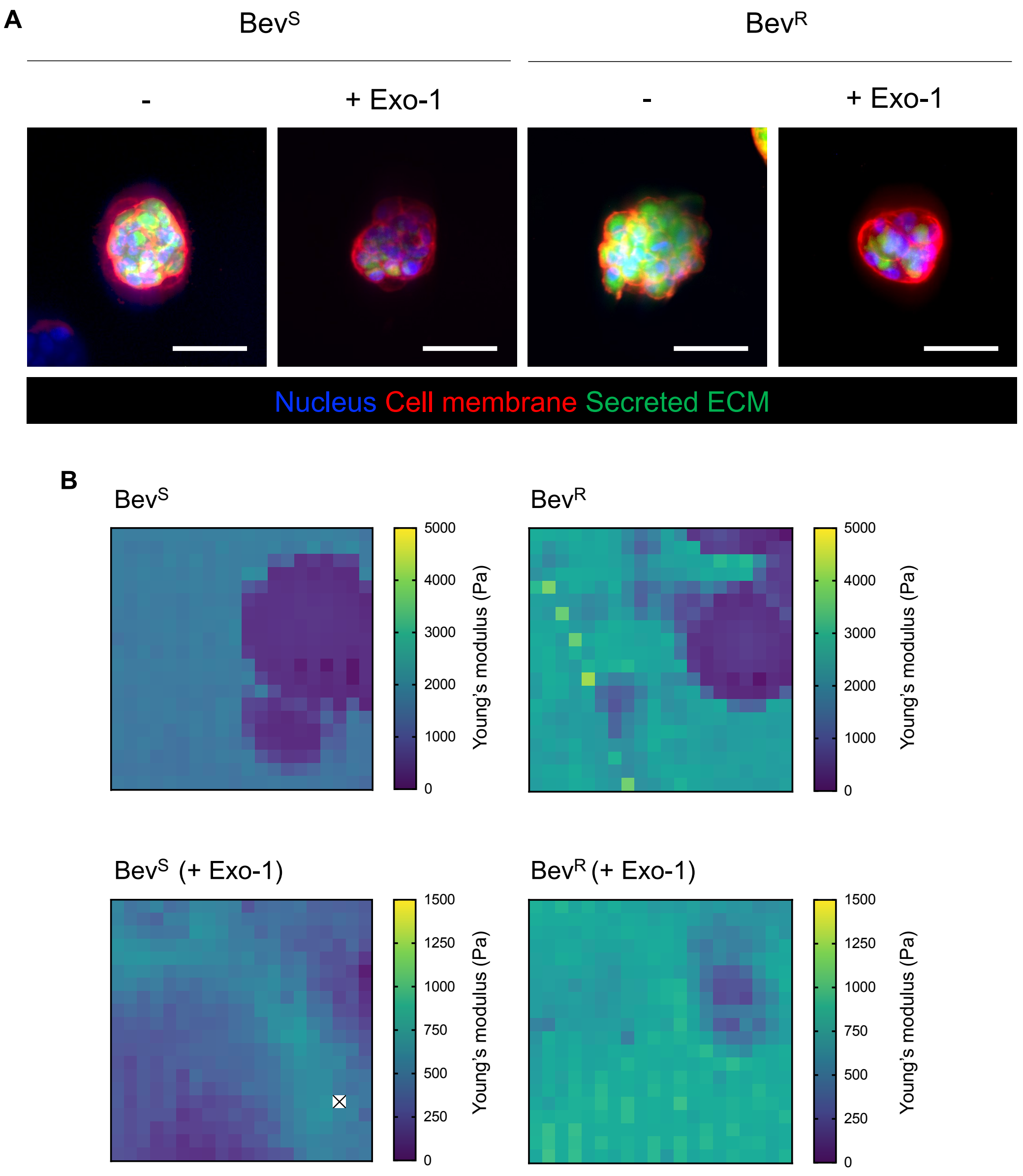


**Fig. S12. Effect of Exo-1 on ECM secretion and matrix stiffness of Bev$^S$ and Bev$^R$ cells-laden HA gels.** (A) Representative fluorescence images of secreted ECM at day 7 with or without Exo-1 treatment. Scale bar: 50 µm. (B) Representative force maps of HA/RGD- gels encapsulating Bev$^S$ and Bev$^R$ cells with/without Exo-1 treatment.

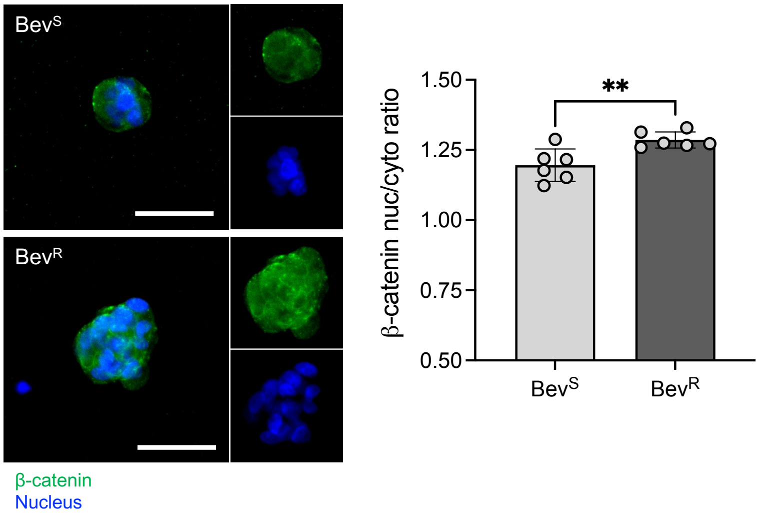


**Fig. S13. β-catenin nuclear localization of Bev$^S$ and Bev$^R$ cells in HA/RGD- gels.** Representative images (left) and quantification (right) of immunofluorescence staining for β-catenin of Bev$^S$ and Bev$^R$ cells in HA/RGD- gels after 7 days. Scale bar: 50 µm. ($n = 6$) Statistical significance was analyzed using an unpaired two-sided Student's t-test. **$p < 0.01$.

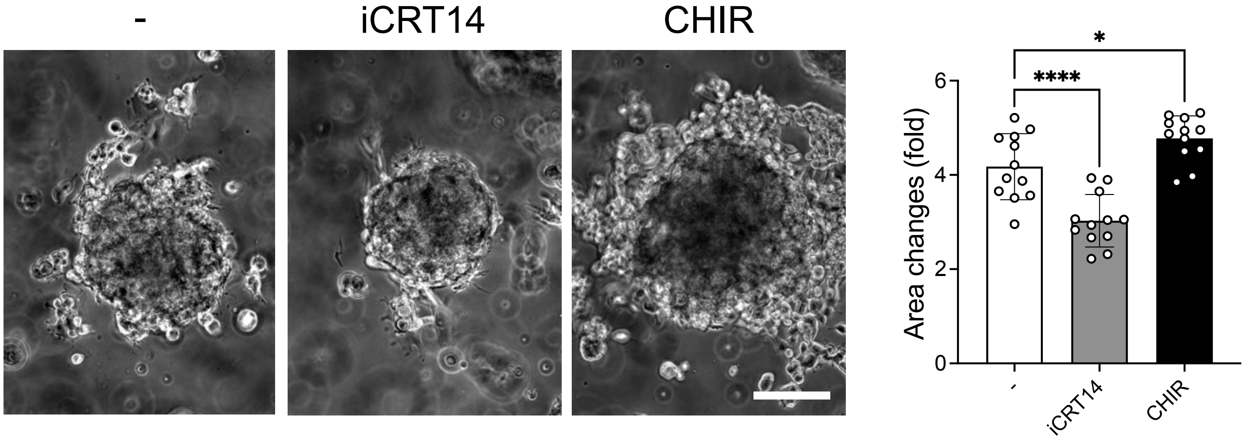


**Fig. S14. Effect of inhibition of β-catenin signaling on GBM tumorsphere invasion.** Representative images (left) and quantification (right) of 3D invasion at day 4 of $Bev^R$ cells with treatment of iCRT14, an inhibitor of β-catenin-TCF/LEF interaction, and CHIR, a highly potent and specific glycogen synthase kinase-3 (GSK3) inhibitor that potentiates β-catenin activity. ($n = 12$) Statistical significance was analyzed using one-way ANOVA followed by Tukey's multiple comparisons test. ****$p < 0.0001$, *$p < 0.05$. Scale bar: 50 μm.

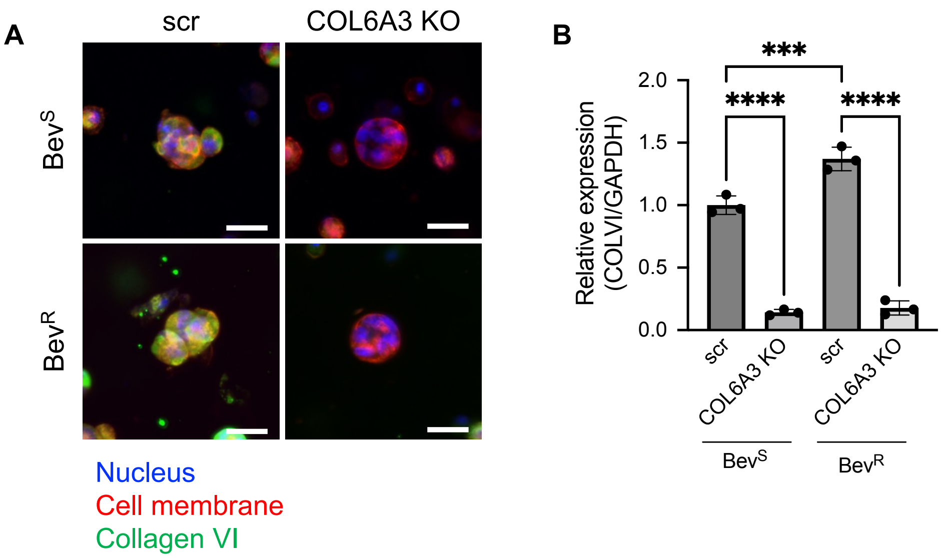


**Fig. S15. COL6A3 knockout (KO) of Bev$^S$ and Bev$^R$ cells using CRISPR gene editing.** (**A**) Representative fluorescence images of collagen VI from non-targeted and COL6A3 gene-deleted Bev$^S$ and Bev$^R$ cells within 3D HA/RGD- at day 7. Scale bar: 30 µm. (**B**) collagen VI mRNA levels in non-targeting and COL6A3 knockout Bev$^S$ and Bev$^R$ cells within 3D HA/RGD- gels. (n = 3) Statistical significance was analyzed using a two-way ANOVA with Tukey's multiple comparisons test. ****$p < 0.0001$, ***$p < 0.001$.

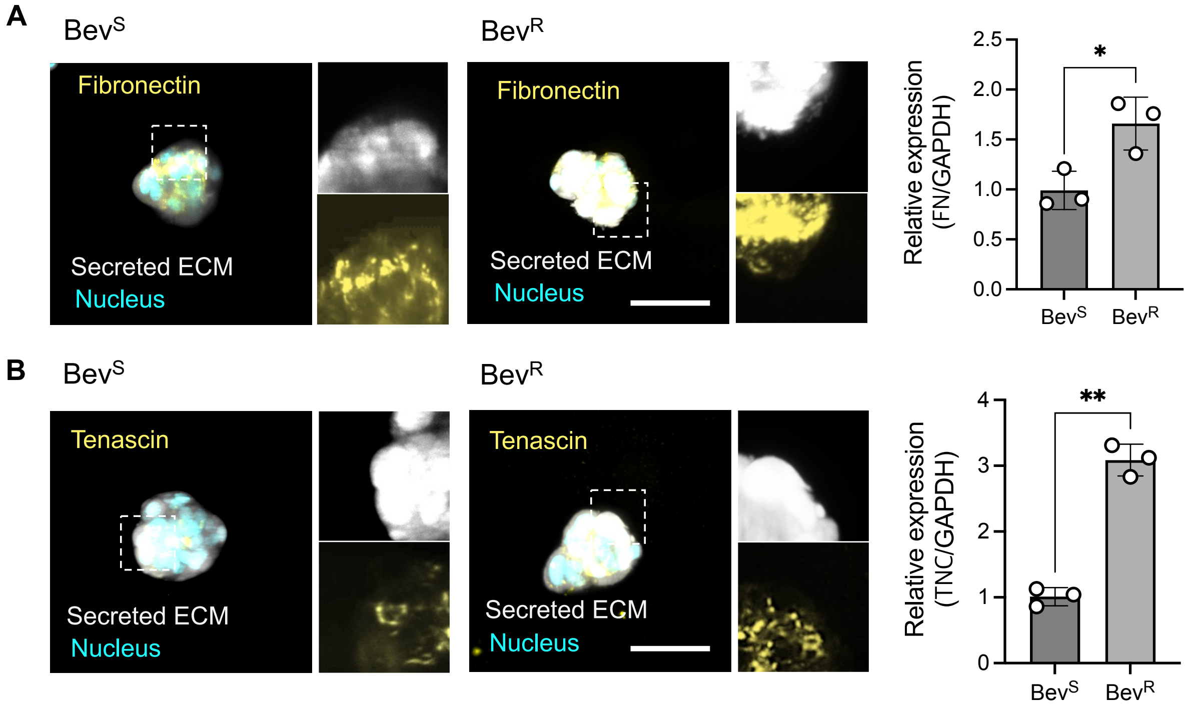


**Fig. S16. Fibronectin and Tenascin secreted by Bev$^S$ and Bev$^R$ cells within 3D HA/RGD- gels.** (**A**) Representative fluorescence images of secreted ECM and fibronectin from Bev$^S$ and Bev$^R$ cells within 3D HA/RGD- gels at day 7 (left). Scale bar: 50 µm. mRNA expression levels of fibronectin in Bev$^S$ and Bev$^R$ cells within 3D HA/RGD- gels. (n = 3) (**B**) Representative fluorescence images of secreted ECM and tenascin from Bev$^S$ and Bev$^R$ cells within 3D HA/RGD- gels at day 7 (left). Scale bar: 50 µm. mRNA expression levels of tenascin in Bev$^S$ and Bev$^R$ cells within 3D HA/RGD- gels. (n = 3) Statistical significance was analyzed using an unpaired two-sided Student's t-test. $^{**}p < 0.01$, $^{*}p < 0.05$.

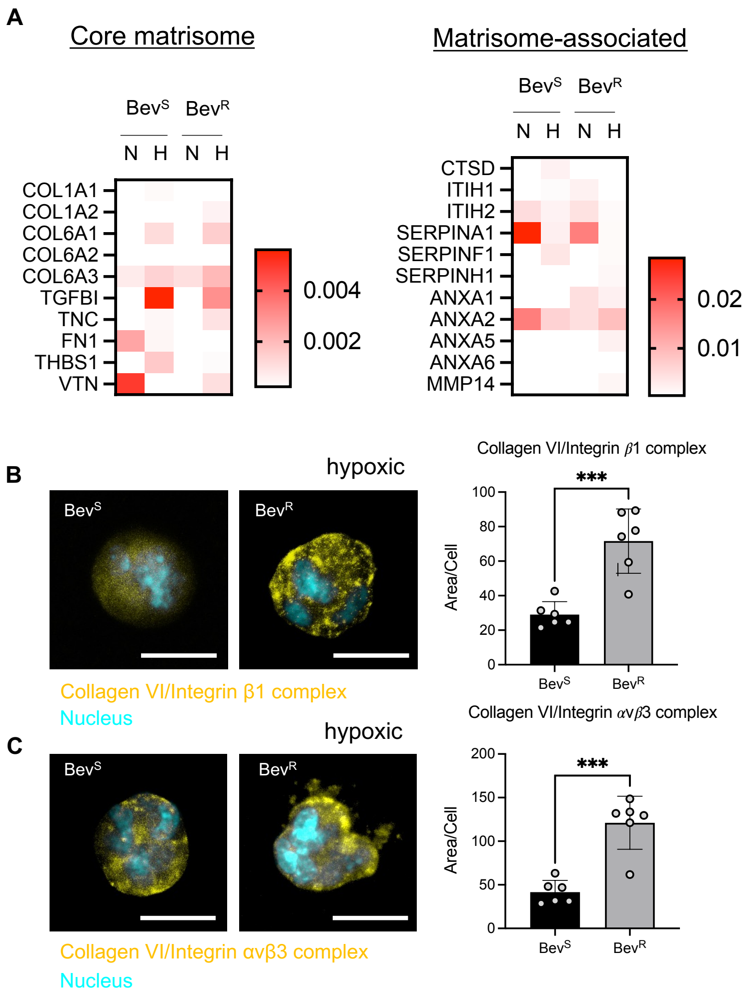


**Fig. S17. Changes of ECM composition and collagen VI-integrin colocalization in Bev$^S$ and Bev$^R$ cells within 3D HA/RGD- hydrogels under normoxic (20%) and hypoxic (1%) conditions.** (**A**) Heatmaps of matrisome proteins secreted by Bev$^S$ and Bev$^R$ cells within 3D HA/RGD- gels under normoxic (20%) and hypoxic (1%) conditions. N: normoxic; H: hypoxic. (**B**, **C**) Proximity ligation assay (PLA) to investigate interactions between secreted collagen VI and integrins under hypoxic conditions. (**B**) Representative images (left) and quantification (right) of collagen VI interactions with integrin β1 in Bev$^S$ and Bev$^R$ cells within 3D HA/RGD- gels under hypoxic conditions. Scale bar: 50 µm. (n = 6) (**C**) Representative images (left) and quantification (right) of collagen VI interactions with integrin avβ3 in Bev$^S$ and Bev$^R$ cells within 3D HA/RGD- gels under hypoxic conditions. Scale bar: 50 µm. (n = 6) Statistical significance was analyzed using an unpaired two-sided Student's t-test. ***$p < 0.001$.

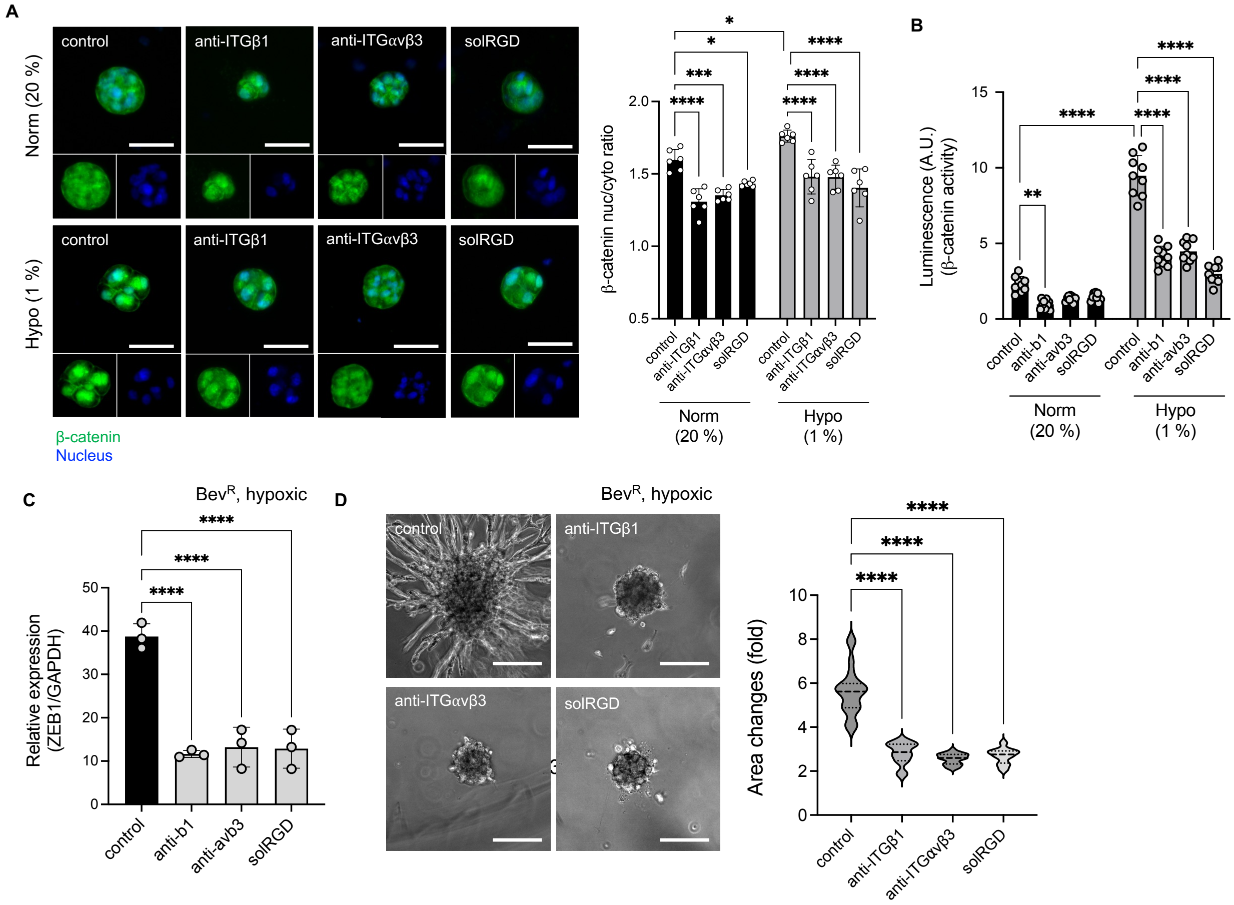


**Fig. S18**. **Effect of inhibition of integrin binding on β-catenin signaling, ZEB1 expression, and invasion.** (**A**) Immunostaining of β-catenin in Bev$^R$ cells in HA/RGD- after 7 days under the normoxic and hypoxic condition of no treatment (control), or treatment with monoclonal antibodies against integrin β1 (anti-ITGβ1), integrin avβ3 (anti-ITG avβ3), or soluble RGD (solRGD) (left). Scale bar: 50 µm. Quantification of β-catenin nuclear localization in Bev$^R$ cells encapsulated within HA/RGD- gels in response to inhibition of integrin binding (right) under normoxic and hypoxic conditions. (n = 6) (**B**) Luciferase assay for assessing β-catenin activity for Bev$^R$ cells encapsulated within HA/RGD- gels in response to inhibition of integrin binding under normoxic and hypoxic conditions. (n = 9) (**C**) ZEB1 gene expression following perturbation of integrin binding in Bev$^R$ cells under hypoxic conditions. (n = 3) (**D**) Representative images (left) and quantification (right) of 3D sphere invasion assay of Bev$^R$ cells under hypoxia in response to perturbation of integrin binding at day 4. Scale bar: 100 µm. (n = 21 to 24) Statistical significance was analyzed using a one-way or two-way ANOVA with Tukey's multiple comparisons test. ****$p < 0.0001$, ***$p < 0.001$, **$p < 0.01$, *$p < 0.05$.

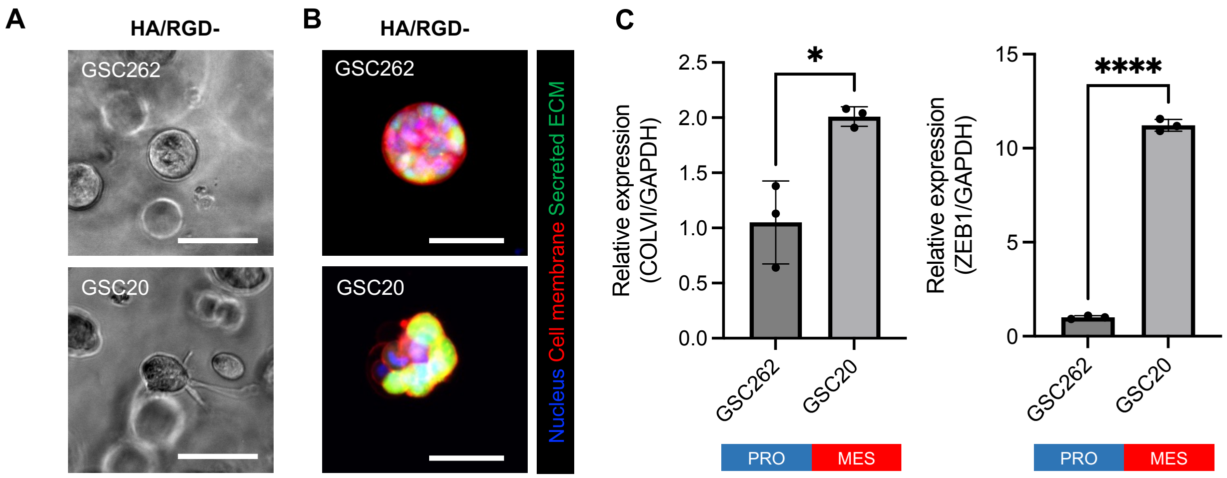


**Fig. S19. Patient-derived GBM stem cells (GSCs) encapsulated within 3D HA/RGD- gels.** (**A**) Representative images of GSCs encapsulated within 3D HA/RGD- hydrogels after 7 days. GSC262: proneural subtype, GSC20: mesenchymal subtype. Scale bar: 100 µm. (**B**) Representative images of fluorescently labeled secreted ECM by GSCs within 3D HA/RGD- gels. Scale bar: 50 µm. (**C**) COLVI gene expression and (**D**) ZEB1 gene expression by GSC262 and GSC20 within 3D HA/RGD- gels. (n = 3) Statistical significance was analyzed using an unpaired two-sided Student's t-test. ****$p < 0.0001$, *$p < 0.05$.
